# Extreme climatic events but not environmental heterogeneity shape within-population genetic variation in maritime pine

**DOI:** 10.1101/2021.08.17.456636

**Authors:** Juliette Archambeau, Marta Benito Garzón, Marina de Miguel Vega, Benjamin Brachi, Frédéric Barraquand, Santiago C. González-Martínez

## Abstract

How evolutionary forces interact to maintain quantitative genetic variation within populations has been a matter of extensive theoretical debates. While mutation and migration increase genetic variation, natural selection and genetic drift are expected to deplete it. To date, levels of genetic variation observed in natural populations are hard to predict without accounting for other processes, such as balancing selection in heterogeneous environments. We aimed to empirically test three hypotheses: (i) admixed populations have higher quantitative genetic variation due to introgression from other gene pools, (ii) quantitative genetic variation is lower in populations from harsher environments (i.e. experiencing stronger selection), and (iii) quantitative genetic variation is higher in populations from spatially heterogeneous environments. We used phenotypic measurements of five growth, phenological and functional traits from three clonal common gardens, consisting of 523 clones from 33 populations of maritime pine (*Pinus pinaster* Aiton). Populations from harsher climates (mainly colder areas) showed lower genetic variation for height in the three common gardens. Surprisingly, we did not find any association between within-population genetic variation and environmental heterogeneity or population admixture for any trait. Our results suggest a predominant role of natural selection in driving within-population genetic variation, and therefore indirectly their adaptive potential.

## 1 Introduction

Most complex traits show substantial heritable variation in natural populations. How evolutionary forces interact to maintain such variation remains a long-standing dilemma in evolutionary biology and quantitative genetics (Johnson and Barton 2005). While mutation and genetic drift have straightforward roles, generating and eliminating variation respectively, the effect of natural selection is more complicated (Walsh and Lynch 2018). Stabilizing selection, i.e. the selection of intermediate phenotypes, is often strong in natural populations (Hereford et al. 2004). This type of selection is expected to deplete genetic variation (Fisher 1930), either directly on the focal trait or indirectly via pleiotropic effects (Johnson and Barton 2005). Theoretical models based on the balance between mutation, drift and stabilizing selection support this idea, but they suggest lower fitness heritability values than those generally observed in empirical studies (Johnson and Barton 2005). Balancing selection encompasses various evolutionary processes that can maintain greater than neutral genetic variation within populations (Mitchell-Olds et al. 2007). The most widely studied of these processes are heterozygote advantage, frequency-dependent selection, e.g. in disease resistance or self-incompatibility systems (Bergelson et al. 2001, Charlesworth et al. 2005), and temporally or spatially fluctuating selection pressures (Felsenstein 1976). The maintenance of stable polymorphism in spatially heterogeneous environments was first theorized by Levene’s archetypal model (1953), under the assumptions of random mating within generations and soft selection. Since then, a large corpus of single-locus and polygenic models, most often deterministic, have generally concluded that genetic polymorphisms can only be maintained under restrictive conditions (Spichtig and Kawecki 2004, Byers 2005). In this line, McDonald and Yeaman (2018) showed with stochastic individual-based simulations that substantial within-population genetic variation can be maintained in spatially heterogeneous environments at intermediate migration rates, regardless of population size. However, the relative importance of the different evolutionary forces driving within-population genetic variation remains largely unknown.

Long-dating empirical work has addressed the evolutionary processes underlying the maintenance of genetic and discrete-trait polymorphisms (reviewed in Hedrick 1986, 2006), e.g. plant-pathogen interactions (Karasov et al. 2014), antagonistic pleiotropy (Carter and Nguyen 2011), environmental heterogeneity (Chakraborty and Fry 2016), and temporal fluctuations (Bergland et al. 2014). Genomics have allowed the broad application of genome-wide scans for signatures of selection. Overall these scans suggest that many loci are under adaptive directional selection (Barreiro et al. 2008, Fu and Akey 2013) and that the proportion of genetic polymorphisms maintained by environmental heterogeneity tends to be low (Hedrick 2006). However those scans typically have low power to detect signatures of balancing selection or local adaptation (Fijarczyk and Babik 2015). Far fewer empirical studies have focused on assessing the distribution and extent of the quantitative genetic variation within populations, and its underlying causes (Lynch and Walsh 1998). Traits more closely related to fitness, such as life-history traits, have generally higher additive genetic variance, but lower heritabilities, than morphometric traits (Price and Schluter 1991, Houle 1992, Kruuk et al. 2000). The hypothesis that populations evolving under strong selection pressures display lower levels of genetic variation has been supported in experimentally evolving quail populations under unfavorable vs favorable treatments (Marks 1978), in controlled experiments (Colautti et al. 2010; but see Merilä et al. 2004, Stock et al. 2014), in natural populations of *Drosophila birchii* subject to climatic selection (but see *D. bennata* and *D. serrata*; van Heerwaarden et al. 2009) and in some natural populations of great tits subject to varying levels of food availability (Charmantier et al. 2004). Higher genetic variation in populations evolving under spatially varying selection pressures is supported by experimental evolution of *Drosophila* populations (Mackay 1981, Huang et al. 2015; but not Yeaman et al. 2010) and in forest trees evaluated in common gardens (Yeaman and Jarvis 2006). The lack of general trends from these empirical studies can be explained by method-specific pitfalls to accurately estimate quantitative genetic variation, e.g. the genetic and environmental variances are hard to disentangle in the wild, and when estimated in common gardens, their environment-dependent nature does not allow for wide generalization of estimates (Hoffmann and Parsons 1991, Merilä et al. 2001, Charmantier et al. 2004). In addition, gene flow has been hypothesized to have either a positive effect on the adaptive potential, by increasing standing genetic variation, or a negative effect via gene swamping (Kremer et al. 2012, Tigano and Friesen 2016), which may depend on the spatial scale considered (Bridle et al. 2009).

Forest trees have specific life-history traits and genomic features making them interesting model species in population and quantitative genetic studies (Petit and Hampe 2006, Savolainen et al. 2007). Compared to crop species, they remain largely undomesticated (Neale and Savolainen 2004). Most forest trees are outcrossing, have high lifetime reproductive output and long generation times. They often display important gene flow among populations through long-distance pollen dispersal (Kremer et al. 2012). They show slow rates of macroevolution (i.e. low nucleotide substitution rates and low speciation rates; Petit and Hampe 2006), generally have large effective population sizes, with distributions often covering a wide range of environmental conditions (Alberto et al. 2013). Extensive work has revealed strong clines at large geographical scales in the population-specific mean values of phenotypic traits (reviewed in Savolainen et al. 2007, Benito Garzón et al. 2019), e.g. phenological traits with latitude or altitude (Alberto et al. 2011, Thibault et al. 2020) or height growth with cold hardiness (Rehfeldt et al. 1999, Leites et al. 2012). Genetic differentiation at microgeographic spatial scales has also been repeatedly observed (reviewed in Linhart and Grant 1996, Jump and Peñuelas 2005, Scotti et al. 2016), suggesting rapid rates of microevolution (Petit et al. 2004, Petit and Hampe 2006). Possible explanations include the fact that forest trees have high levels of genetic diversity and that most of their quantitative and neutral genetic variation is within populations (Hamrick 2004). To our knowledge, only two empirical studies investigated the potential causes underlying the maintenance of quantitative trait variation within forest tree populations. Yeaman and Jarvis (2006) showed that 20% of growth genetic variation in lodgepole pine populations was attributable to regional heterogeneity, suggesting an important role of gene flow and varying selection pressures. In the neotropical oak *Q. oleoides*, Ramírez-Valiente et al. (2019) found lower quantitative genetic variation in harsher environments, but not higher quantitative genetic variation in temporally fluctuating environments. They also suggested only a marginal effect of genetic structure and diversity on the maintenance of within-population genetic variation.

In this study, we aimed to test competing hypotheses regarding the relationship between quantitative genetic variation within maritime pine populations and the potential underlying drivers that maintain this variation. We used phenotypic measurements of growth (height), phenological (bud burst and duration of bud burst) and functional (*δ*^13^C and specific leaf area, SLA) traits from three clonal common gardens, consisting of 522 clones (i.e. genotypes) from 33 populations, spanning all known gene pools in the species (Jaramillo-Correa et al. 2015) and genotyped for 5,165 SNPs. For each trait, we compared Bayesian hierarchical models that estimate the relationship between the total genetic variances within populations and some potential drivers, namely climate’s harshness at the locations of origin of the populations (i.e. drought intensity and severe cold events), environmental heterogeneity in the forested areas surrounding the populations, and the level and origin of admixture in the populations, as estimated with SNP markers. The competing, but not mutually exclusive, hypotheses tested are: i) the most admixed populations have higher quantitative genetic variation due to introgression from other gene pools, and this relationship is proportional to the divergence between sink and source gene pools; ii) quantitative genetic variation is lower in populations that have evolved in harsher environments, as a result of higher selection pressures in these regions; and iii) quantitative genetic variation is higher in populations that have evolved in spatially heterogeneous environments. Importantly, the last two hypotheses require the action of natural selection, while the first does not. Therefore, we expect the last two hypotheses to be mostly supported for fitness-related traits, while the first hypothesis may apply uniformly to all traits. Determining the patterns of within-population quantitative genetic variation across species’ ranges and the relative importance of the evolutionary forces driving the maintenance of such variation is necessary to assess the evolutionary potential of forest tree populations. Empirical studies tackling these questions remain extremely rare in forest trees (but see Yeaman and Jarvis 2006, Ramírez-Valiente et al. 2019), yet they are much needed to anticipate forest tree responses to ongoing global change and therefore develop adaptive management and conservation strategies.

## 2 Materials & Methods

### 2.1 Maritime pine, a forest tree growing in heterogeneous environments

Maritime pine (*Pinus pinaster* Ait., Pinaceae) is a wind-pollinated, outcrossing and long-lived tree species with large ecological and economical importance in western Europe and North Africa. Maritime pine is largely appreciated for its wood, for stabilizing coastal and fossil dunes and, as a keystone species, for supporting biodiversity (Viñas et al. 2016). The distribution of maritime pine natural populations is scattered and covers a wide range of environmental conditions. Several studies have provided evidence of genetic differentiation for adaptive traits in this species, suggesting local adaptation (e.g. González-Martínez et al. 2002, de Miguel et al. 2020). Maritime pine can grow in widely different climates: the dry climate along the northern coasts of the Mediterranean Basin (from Portugal to western Italy), the mountainous climates of south-eastern Spain and Morocco, the wetter climate of the Atlantic region (from the Spanish Iberian region to the western part of France) and the continental climate of central Spain. Maritime pine can also grow on a wide range of substrates, from sandy and acidic soils to more calcareous ones. Maritime pine presents a strong population genetic structure with occasional admixture, suggesting gene flow among gene pools. Six gene pools have been described by previous literature, located in the French Atlantic region, Iberian Atlantic region, central Spain, south-eastern Spain, Corsica and Northern Africa (Fig. 1; Alberto et al. 2013, Jaramillo-Correa et al. 2015). These gene pools probably result from the expansion of different glacial refugia (Bucci et al. 2007, Santos-del-Blanco et al. 2012).

**Figure 1.**
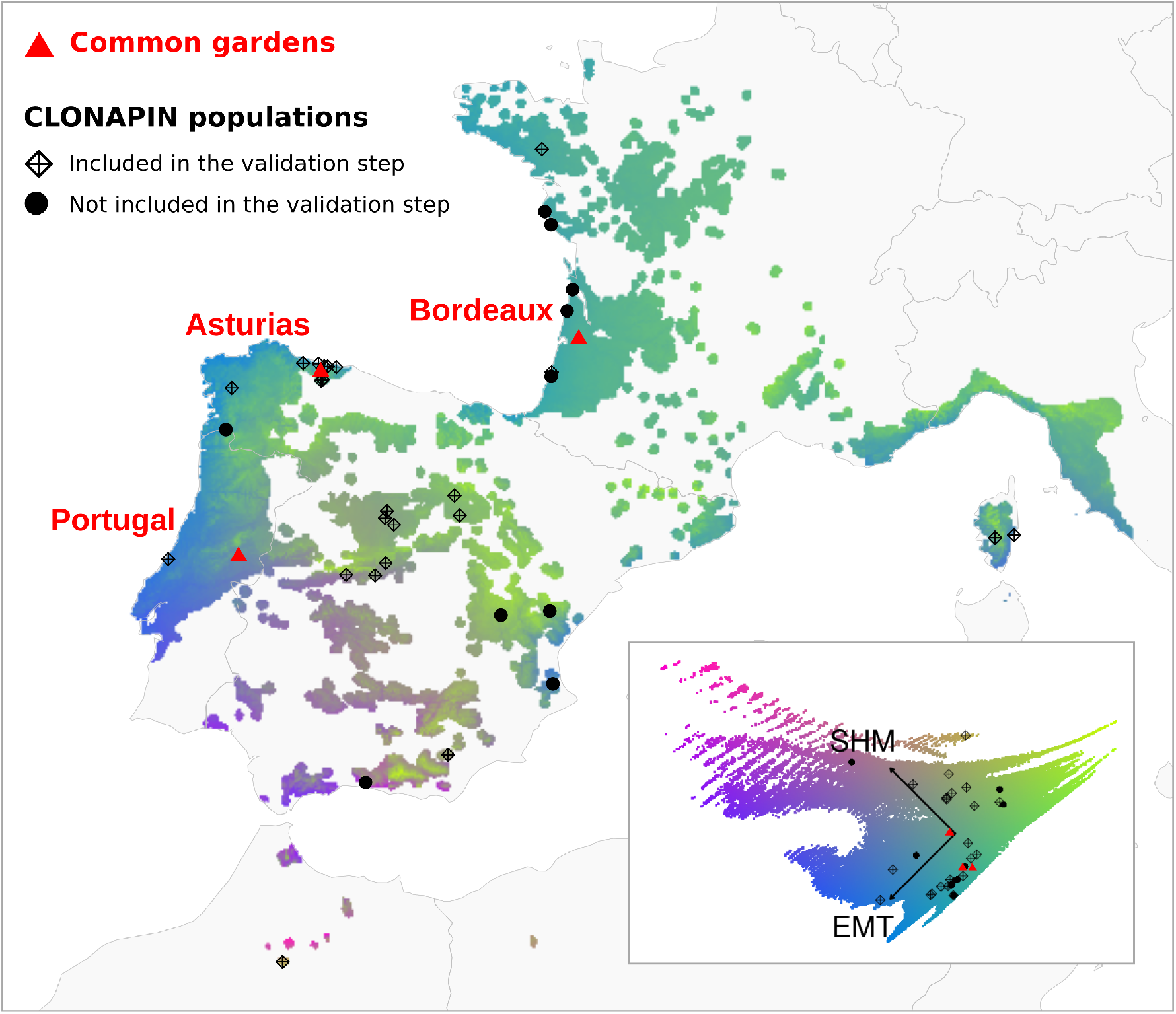
Location of the three common gardens and the 33 populations used in the study. The colors represent the gradients of the extreme minimum temperature (EMT) and summer heat moisture index (SHM) over the period 1901-1950 within the maritime pine range. The climatic gradients were obtained by performing a centered and scaled principal component analysis (shown in the inset on the bottom right) based on EMT and SHM values. The maritime pine distribution combines the EUFORGEN distribution (http://www.euforgen.org/) and 10-km radius areas around the National Forest Inventory plots with maritime pines. However, this remains a rough approximation of the actual distribution of maritime pine and therefore probably includes areas experiencing more intense cold or drought episodes than the climatic range of maritime pine.

### 2.2 Phenotypic data

Phenotypic data was obtained from three clonal common gardens (Table S1 and Fig. 1), planted in 2011 and located in environments considered favorable to maritime pine, as evidenced by the high survival rate at these sites (Table S1). The common gardens of Asturias (Spain, Iberian Atlantic region) and Bordeaux (France, French Atlantic region) have very similar climates, with mild winters, no severe cold events, high annual rainfall and relatively wet summers (Tables S3-S5 and Fig. 1). The common garden of Portugal (planted in Fundão) shows slightly colder winters and lower summer precipitation than in Asturias (Table S4 and Fig. 1). In each of these common gardens, trees belonging to 522 clones (i.e. genotypes) from 33 populations, including the six known gene pools in the species, were planted following a randomized complete block design with 8 blocks, 8 trees per clone and from 2 to 28 clones per population (with an average of 15). To obtain the clones, trees at least 50 m apart were sampled in natural stands, and one seed per tree was planted in a nursery and vegetatively propagated by cuttings (see Rodríguez-Quilón et al. 2016 for details). Clones were therefore considered unrelated.

One growth trait, height, was measured in all common gardens and at different tree ages (Table S1). Two phenology-related traits, the mean bud burst date over four years and the mean duration of bud burst over three years, were measured in Bordeaux and were averaged over several years to suppress differences across years and approximate a normal distribution of their trait values (Table S1). Bud burst corresponds to the date of brachyblast emergence in accumulated degree-days (with base temperature 0°C) from the first day of the year to account for between-year variability in temperature. The duration of bud burst corresponds to the number of degree-days between the beginning of bud elongation and the total elongation of the needles (see Hurel et al. 2019). Last, two functional traits, *δ*^13^C and the specific leaf area (SLA) were measured in Portugal (Table S1). These traits were selected because they showed broad-sense heritabilities that were mostly low but with credibility intervals not crossing zero (*>* 0.08 in de Miguel et al. 2020). For each trait, phenotypic means and variances across populations are shown in section 1.1 of the Supplementary Information. Prior to analyses, some traits were log-transformed to get closer to normality or mean-centered to help model convergence (Table S1).

### 2.3 SNP genotyping and population admixture

The 522 clones planted in the Asturias common garden were genotyped with the Illumina Infinium assay described in Plomion et al. (2016), resulting in 5,165 high-quality polymorphic SNPs. There were on average only 3.3 missing values per genotype (ranging between 0 and 142). For each clone, the proportions of ancestry from each of the six known gene pools were estimated in Jaramillo-Correa et al. (2015) using the Bayesian approach available in Structure (Pritchard et al. 2000), and were then averaged by population. Populations were assigned to the gene pool that contributed more than 50% ancestry and the other gene pools were considered as ‘foreign‘ gene pools. First, we calculated a population admixture score A, as the proportion of ancestry from foreign gene pools (Table S6). Second, we calculated a population admixture score D that considers both the proportion of foreign ancestries and the divergence between the main and foreign gene pools (Table S6). For that, we weighted the proportions of ancestry from foreign gene pools by the sum of the allele frequency divergence of the main and foreign gene pool from the common ancestral one (F_*k*_, which should be numerically similar to F_*ST*_ ; Falush et al. 2003). We developed D considering that some gene pools are more divergent than others and thus may bring higher genetic diversity to an admixed population at the same level of introgression. A and D were highly correlated (Pearson correlation coefficient of 0.91; Table S8). We also calculated a score D_*fst*_ by weighting the proportions of ancestry from foreign gene pools by the pairwise F_*ST*_ between the main and foreign gene pools (Table S8). D_*fst*_ was highly correlated to A and D (Pearson correlation coefficients of 0.91 and 0.96, respectively) and we therefore did not keep it in the following analyses.

### 2.4 Population-specific environmental heterogeneity and climate harshness indexes

To describe the climate under which the populations have evolved, we used the climatic variables at 1-km resolution and averaged over the period 1901-1950 from the *ClimateEU* database (Marchi et al. 2020). Topographic data were generated from NASA’s Shuttle Radar Topography Mission (SRTM) at 90-m resolution and then aggregated at 1-km resolution. We used the SAGA v 2.3.1 (Conrad et al. 2015) to calculate the topographic ruggedness index (TRI) which quantifies the terrain heterogeneity, i.e. differences in elevation between adjacent cells (Riley et al. 1999). Soil variables were extracted from the European Soil Database at 1-km resolution (Hiederer et al. 2013). All environmental variables used are listed in Table S7 and were mean-centered and divided by their standard deviation prior to analyses.

To calculate the environmental heterogeneity around each population location, we extracted raster cell values of the climatic, topographic and soil variables within a 20-km radius around each population location, and kept only raster cells that fell within forested areas, to avoid including environmental data from non-suitable areas (e.g. lakes, mountain peaks; section 1.3.2 of the Supplementary Information). We then performed a principal component analysis (PCA) on the raster cell values and extracted the PC1 and PC2 scores of each cell, accounting for 45.2% and 34.1% of the variance, respectively (Fig. S10). To obtain the four indexes of environmental heterogeneity, we calculated the variances of the PC1 and PC2 scores in a 20-km and 1.6-km radius around each population location. The environmental heterogeneity indexes were only very weakly correlated (Pearson correlation coefficients lower than 0.36) with the number of forested cells (i.e. the area considered to calculate the indexes), ensuring that the estimated effects of environmental heterogeneity in further analyses were not due to the area per se (Triantis et al. 2003, Stein et al. 2014).

To describe the climate harshness at each population location, we used a drought index (the summer heat moisture index averaged over the period 1901-1950, SHM, Table S7) and an index related to severe cold events (the inverse of the extreme minimum temperature during the period 1901-1950, invEMT, Table S7). These two indexes were selected as maritime pine shows local adaptation patterns associated with cold tolerance (Grivet et al. 2011) and because detecting changes in the within-population genetic variation along a drought gradient would be key to anticipate tree population responses to ongoing climate change.

### 2.5 Bayesian statistical modeling

We modeled the eight phenotypic traits with the same Bayesian statistical model, in which we estimate the linear relationship between the within-population genetic variance and each of the potential drivers successively (i.e. one model per driver): the two admixture scores, the four environmental heterogeneity indexes and the two climate harshness indexes. Each trait *y* followed a normal distribution (Fig. S1), such as:

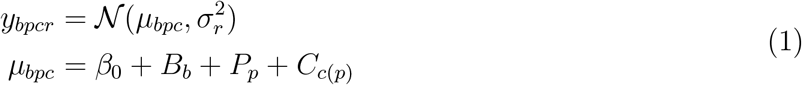

where 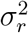 is the residual variance, *β*_0_ the global intercept, and *B*_*b*_, *P*_*p*_ and *C*_*c*(*p*)_ are the block, population and clone (nested within population) varying intercepts, which are drawn from a common distribution, such as:

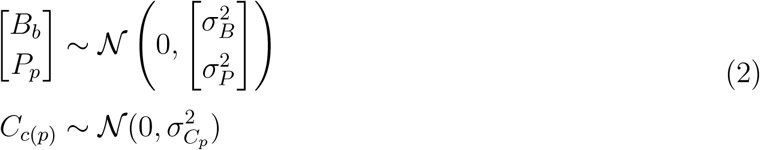

where 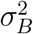 and 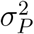 are the variance among blocks and populations and 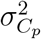 are the population-specific variances among clones (i.e. the within-population genetic variation). To estimate the association between 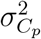 and its potential underlying drivers, we expressed 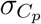 as follows:

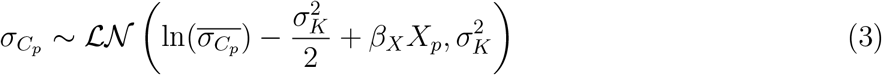

where 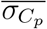 is the mean of the population-specific standard deviation among clones 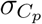 and *X*_*p*_ is the potential driver considered (see section 2 in the Supplementary Information for more details).

To test the accuracy of the model estimates for 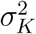 and *β*_*X*_, we simulated data based on two traits (height in Portugal and Bordeaux at 20 and 25-month old, respectively). For each trait, we ran 100 simulations and extracted the mean standard error and bias error of the estimates and the coverage of the 80% and 95% credible intervals.

Model specification and fit were performed using the Stan probabilistic programming language (Carpenter et al. 2017), based on the no-U-turn sampler algorithm. Models were run with four chains and between 2,500 iterations per chain depending on the models (including 1,250 warm-up samples not used for the inference). All analyses were undertaken in R version 3.6.3 (R Core Team 2020) and scripts are available at https://github.com/JulietteArchambeau/H2Pinpin.

### 2.6 Validation step on independent data

To validate our results for height, we used an independent dataset provided by Ricardo Alia in which 23 populations shared with the CLONAPIN network were planted in a progeny test near Asturias (thus in a similar environment). As the progeny test is based on families, we were able to estimate the additive genetic variance within populations. We applied the same model as in our study (replacing clones by families) to height measurements when the trees were 3 and 6-year old (see section 7 of the Supplementary Information for more details).

## 3 Results

In the data simulation, 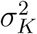 (the standard deviation of the logarithm of the within-population genetic variation) and *β*_*X*_ (the coefficient of the potential drivers of the within-population genetic variation) were properly estimated by the models (Table S9 and S10). Across 100 simulations, the mean standard error was around 0.066 for 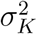 and 0.054 for *β*_*X*_, the mean bias error was around 0.018 for 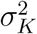 and -0.004 for *β*_*X*_, the coverage of the 80% credible interval was around 93% for 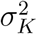 and 80% for *β*_*X*_, and the coverage of the 95% credible interval was around 98% for 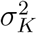 and 96% for *β*_*X*_ (Table S9 and S10). These simulations therefore showed that, under the assumption that the statistical model reflects the processes at work, our model displayed a satisfactory accuracy to be used in the following analyses.

The proportion of variance explained by the models (i.e. the sum of the among-population, among-clone and among-block variances) and the variance partitioning varied broadly across traits (Fig. S11 and section 5.3 in the Supplementary Information). More specifically, the models explained between 40% and 50% of the variance for phenology-related traits, between 30% and 40% for functional traits, and from 20% for height in Portugal to almost 60% for height in Bordeaux at 85-month old (Fig. S11). Residual variance explained most of the variance for all traits, except for height in Bordeaux at 85-month old, where 40% of the variance came from variation among populations, 40% from residuals and the remaining 20% from variation among clones (Fig. S18). Variation among populations was higher than variation among clones for height and *δ*_13_C (Figs. S14, S16, S18, S20 and S28), but not for SLA and phenology-related traits (Figs. S26, S22 and S24).

Environmental heterogeneity indexes and population admixture scores were not associated with within-population genetic variation for any trait (Figs. 2 and S12). In contrast, we found a consistent negative association with the inverse of the extreme minimum temperature across the three common gardens for height, indicating that populations undergoing severe cold events display less genetic variation (Fig. 2). Interestingly, in the Bordeaux common garden, this negative relationship was found at 25-month old, but not at 85-month old (Fig. 2). A negative association with the summer heat moisture index was also detected for height in Asturias, and less markedly but still with a high probability in Bordeaux at 25-month old (Fig. 2). Holding all other parameters constant, a one-standard deviation increase in the inverse of the extreme minimum temperature was associated, on average, with a 32.6%, 21.6% and 17.9% decrease of 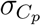 for height in Portugal, Bordeaux at 25-month old and Asturias, respectively. Similarly, a one-standard deviation increase in the summer heat moisture index was associated, on average, with 15.6% and 23.8% decrease of 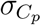 for height in Bordeaux at 25-month old and Asturias, respectively (see details of the calculation in the section 4 of the Supplementary Information). Unexpectedly, populations experiencing severe cold events showed higher genetic variation for SLA (Fig. 2). Within-population genetic variation was not correlated with the number of clones per population for any trait (maximum Pearson correlation coefficient = 0.57; Table S11).

**Figure 2.**
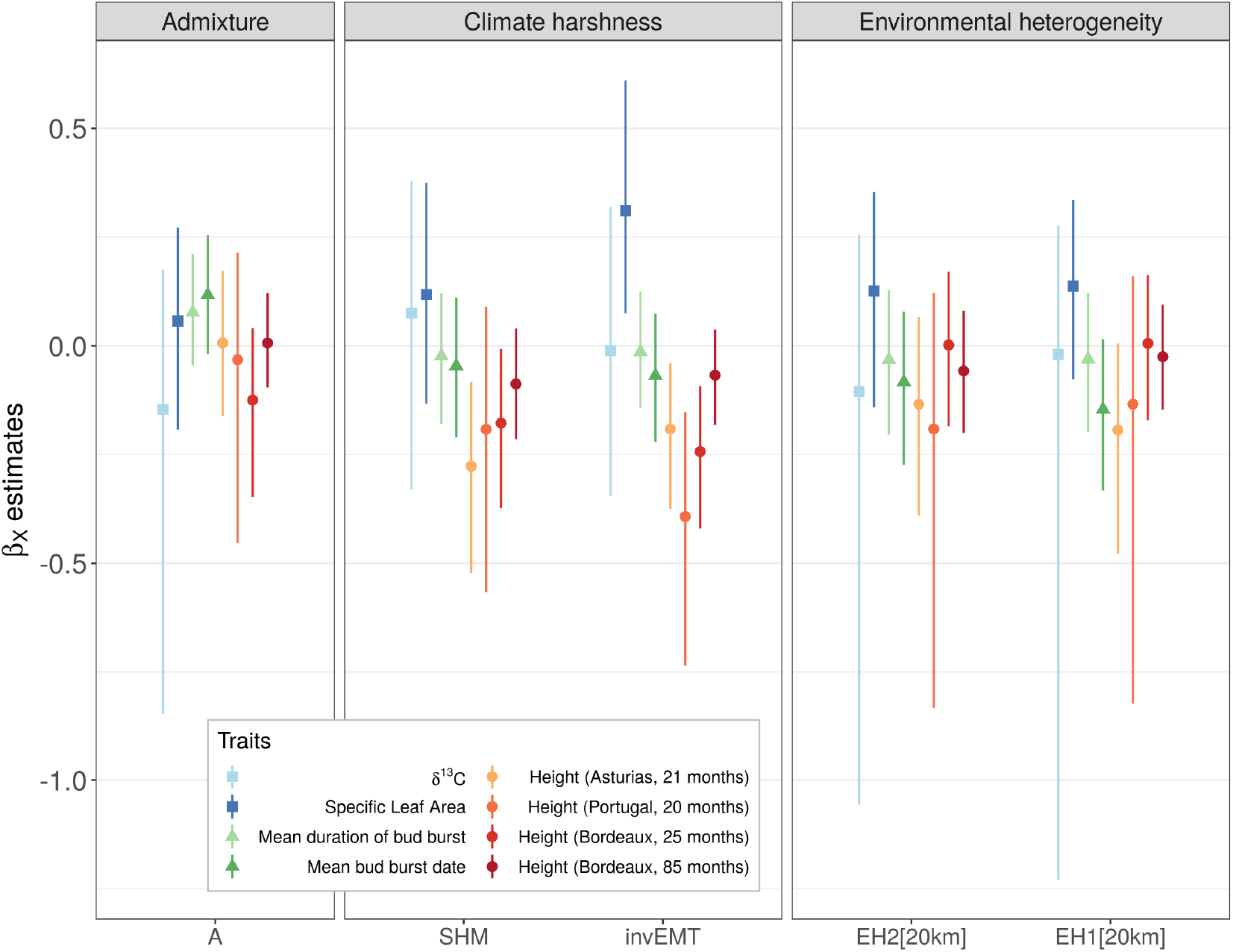
Median and 95% credible intervals of the *β*_*X*_ posterior distributions. *β*_*X*_ coefficients stand for the association between the within-population genetic variation and its potential underlying drivers on the x-axis: the inverse of the extreme minimum temperature during the studied period (invEMT), the summer heat moisture index (SHM), an admixture score (A), the environmental heterogeneity in a 20-km radius around the population location (EH1[20km] and EH2[20km]) calculated based on the projection of the PC1 and PC2 scores. Colors stand for the different traits under study and the shapes for the different types of traits, i.e. functional traits (squares), phenology-related traits (triangles) and height (circles).

Importantly, in the validation analysis, we also found a negative association between the inverse of the extreme minimum temperature and the within-population additive genetic variation for height at 3-year old, but not at 6-year old, and we did not find any association with the other potential drivers (Fig. 3).

**Figure 3.**
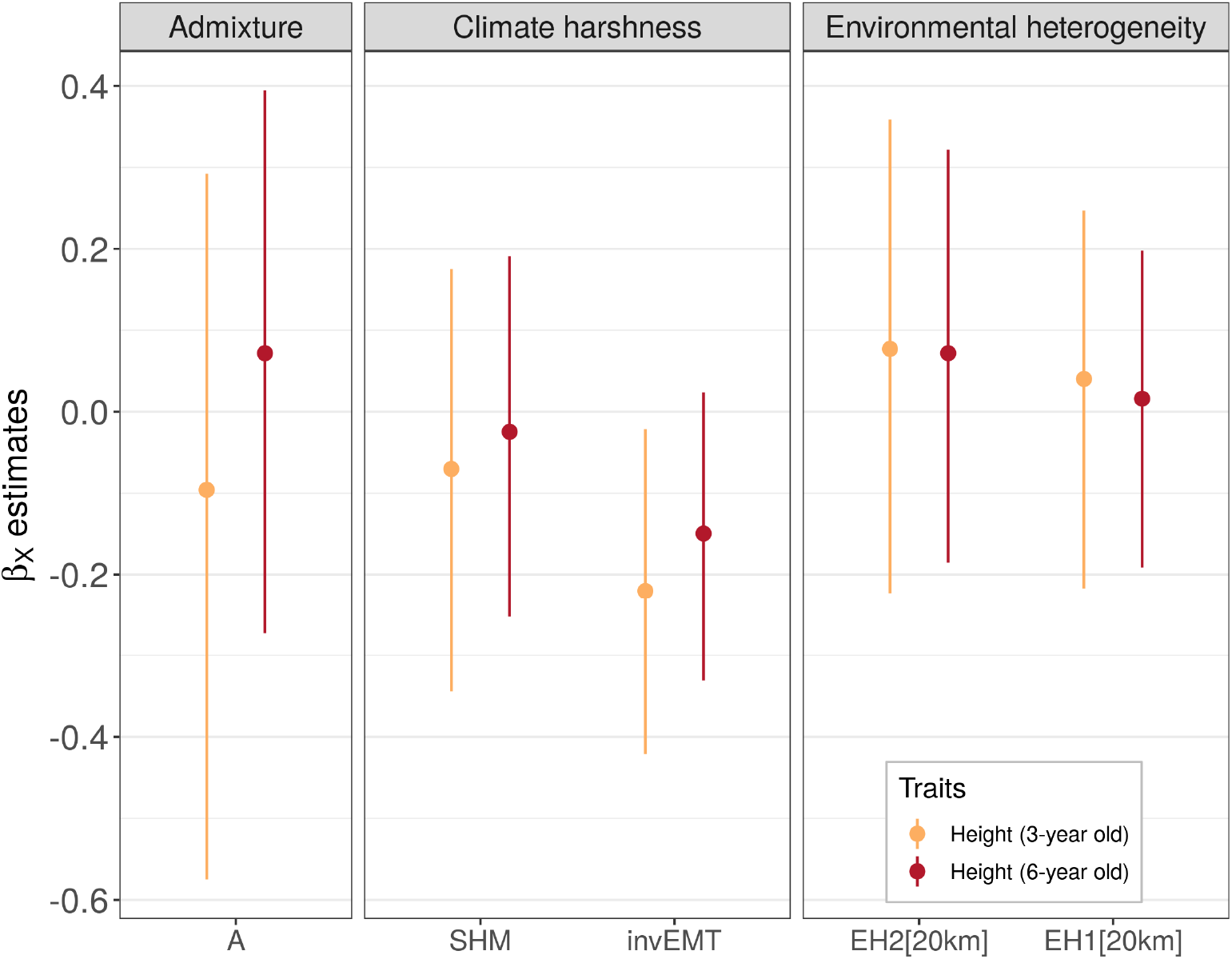
Validation step using independent height measurements from a common garden near Asturias. Median and 95% credible intervals of the *β*_*X*_ posterior distributions are shown. In the validation analysis, *β*_*X*_ coefficients stand for the association between the within-population additive genetic variation and its potential underlying drivers on the x-axis. A description of the drivers can be found in the legend of Fig. 2.

## 4 Discussion

How quantitative genetic variation is maintained within populations remains a long-standing open question that has been extensively explored in theoretical work but lacks empirical evidence to date (Johnson and Barton 2005). Our study suggests that genetic variation for height in maritime pine is lower in populations exposed to severe cold events, thus supporting the hypothesis that quantitative genetic variation in fitness-related traits is lower in populations under strong selection (Fisher 1930). Across all traits studied, we did not find higher genetic variation in populations located in heterogeneous landscapes, which goes against the predictions of some theoretical models (McDonald and Yeaman 2018, Walsh and Lynch 2018) and an empirical study in lodgepole pine (Yeaman and Jarvis 2006). Admixed populations did not show higher genetic variation, suggesting that the observed patterns are not confounded by gene flow between distinct gene pools increasing genetic variation. Empirically-based detection of the footprints of natural selection on within-population genetic variation is much needed to understand how populations are adapted to their current environments and will evolve under changing conditions.

### 4.1 Severe cold events may decrease within-population genetic variation

Height genetic variation was lower in populations experiencing harsher climates, especially severe cold events (invEMT; Fig. 2). This result supports the hypothesis that strong stabilizing selection in harsh environments depletes quantitative genetic variation within populations (Fisher 1930) and echoes similar results in another forest tree, *Quercus oleoides*. For this Mesoamerican white oak species, Ramírez-Valiente et al. (2019) found lower genetic variation averaged over functional and growth traits in populations experiencing low precipitation and high temperatures during the dry season. The importance of severe cold events as a driver of height genetic variation in maritime pine is supported by the association between candidate-gene allele frequency and temperature gradients (Grivet et al. 2011, Jaramillo-Correa et al. 2015), suggesting a major role of minimum temperatures in the species adaptive evolution. Indeed, lower genetic variation in areas subject to cold events may enhance adaptation to local conditions, but it may also hamper the adaptive potential of populations under new climates. Noticeably, severe cold events were highly correlated with altitude in our study (Pearson’s correlation of 0.9), and adaptation patterns along altitudinal gradients are common in forest trees (e.g. Kurt et al. 2012). Therefore, we cannot exclude that the association between height genetic variation and severe cold events is triggered by more complex environmental factors typical of high altitude conditions (e.g. reduced vapor pressure deficit, higher maximum solar radiation; Körner 1995). Estimating selection strength directly in natural populations, as in Bontemps et al. (2016), would be highly valuable, albeit challenging in forest trees.

Within-population genetic variation was unlikely to be influenced by demographic history and gene flow across gene pools, as we did not find any association between within-population genetic variation for height (and other traits) and population admixture indexes (Fig. 2). Another potential scenario that may explain our results is related to the joint effect of environment on the short-term expression of quantitative genetic variation and the strength of natural selection in the novel environments of the common gardens (Hoffmann and Merilä 1999, Wood and Brodie 2016; see examples for natural populations in Wilson et al. 2006 for wild sheep and Husby et al. 2011 for great tits). Noticeably, the expression of hidden genetic variation in novel environments (i.e. ‘cryptic genetic variation’; Schlichting 2008) may be as large as the genetic variation resulting from the long-term divergent evolution of populations (Wood and Brodie 2015). However, in our study, this scenario is unlikely as the negative associations between height genetic variation and climate harshness were consistent across the three common gardens (i.e. across distinct environmental conditions; partially reflected in Fig. 1; see also Tables S3-S5), which suggests that the lower height genetic variation in populations from harsher climates is independent from the environmental conditions in the common gardens and thus likely to be intrinsic to the populations. Last, the sampled populations may not fully cover the climatic range of maritime pine (Fig. 1), which reduces our ability to detect an association between some climatic drivers and within-population genetic variation, and therefore may explain why no association was detected for the summer heat moisture index (SHM).

Most importantly, the validation analysis provided independent evidence that additive within-population genetic variation for height was lower in populations experiencing extreme cold events for young trees but not for older trees (Fig. 3). This supports the robustness of our study and suggests that our results were unlikely to be biased by considering the total variance instead of the additive one, which was somehow expected as two previous studies in maritime pine found low non-additive effects for growth (Gaspar et al. 2013), and height and diameter (Lepoittevin et al. 2011).

With respect to specific leaf area (SLA), where only a single common garden (i.e. a single environment) was assessed, cryptic genetic variation (as defined above) may indeed underlie the higher genetic variation found in populations experiencing severe cold events. A study in maritime pine suggests that SLA depends strongly on environmental conditions (Alía et al. 2014), which is supported in our study by its weak genetic variation in the Portugal common garden (less than 10% of the phenotypic variance is explained by the population or clone effects), with a large part of the variance associated with the block effect (Fig. S26). Cryptic genetic variation is more likely to be expressed when the differences between original and current environments are large (Paaby and Rockman 2014). Some populations experiencing severe cold events (and high altitude conditions) may therefore have reached the threshold inducing a release of cryptic variation in the Portugal common garden. However, this is not a general pattern as we did not find any association between the climatic transfer distances (i.e. the absolute difference between the climate in the population and the climate in the test site) and the within-population genetic variation for SLA (see section 6 of the Supplementary Information). Replicating SLA measurements in common gardens at high altitude or experiencing extreme cold episodes would be highly valuable to test this hypothesis.

### 4.2 Environmental heterogeneity is not associated with higher genetic variation

Populations from heterogeneous environments did not show higher genetic variation for any trait (Fig. 2), which was also the case for the independent height data from the validation analysis (Fig. 3). This goes against a previous estimate in lodgepole pine suggesting that up to 20% of the genetic variation in growth within populations is explained by environmental heterogeneity (Yeaman and Jarvis 2006). A potential explanation of this discrepancy is the smaller experiment size in our study compared to that of Yeaman and Jarvis (103 populations with an average of 28 planting sites per population). However, in our study, we obtained reasonable credible intervals for most traits (allowing the detection of associations with other drivers) and data simulations suggested that our models have adequate power, rendering this explanation unlikely.

Another explanation is that genetic variation within populations is not affected by the environmental heterogeneity at the regional scale imposed by the 1 *×* 1 km resolution of our climate dataset but at finer spatial scales (also discussed in Yeaman and Jarvis 2006). Indeed, populations can adapt along microgeographic environmental gradients despite the homogenizing effect of gene flow (Richardson et al. 2014), even for forest tree populations with their long-generation times and large effective population sizes (Scotti et al. 2016). However, a correlation between regional and microgeographic environmental heterogeneity across the maritime pine range is very likely: populations showing the highest environmental heterogeneity in our study were located in mountainous areas in which we also expect higher microgeographic variation, e.g. the Cómpeta population (COM) located in the Tejeda and Almijara mountains (southern Spain), the Arenas de San Pedro population (ARN) located in the Sierra de Gredos (central Spain) or the Pineta population (PIE) located close to the Punta di Forchelli (Corsican mountains), while populations with the lowest environmental heterogeneity were located on flat plateaus, e.g. populations from the Landes plateau and the Atlantic coastal regions in France (HOU, MIM, PET, VER, OLO, STJ, PLE), and populations from the central Spain plateau near to Segovia (CUE, COC, CAR). Thus, even if genetic variation was maintained by migration-selection balance at microgeographic scales, we would have been able to detect the effect of environmental heterogeneity at the regional scale. Nevertheless, more studies characterizing adaptation at microgeographic scales are needed to assess the spatial scale of genetic adaptation in maritime pine.

Another explanation of the discrepancy with Yeaman and Jarvis (2006) could be that we used young trees (between 20 and 85-month old) while they used 20-year old trees. Indeed, the processes generating within-population genetic variation might be age-dependent, as shown for climate harshness in Bordeaux, where the association was present when the trees were 25-month old but not in older trees. In forest trees, genetic parameters often vary with age; e.g. heritability generally increases with age until reaching a plateau, especially for height-related traits (Balocchi et al. 1993, Johnson et al. 1997, Sierra-Lucero et al. 2002, Jansson et al. 2003, Kroon et al. 2011), but may also decrease in some cases (Lu and Charrette 2008, Kroon et al. 2011). In maritime pine, an increase in heritability with age was found in Costa and Durel (2011) but not in Kusnandar et al. (1998). To our knowledge, the drivers of heritability changes with age remain unclear. Competition among trees in common gardens might play a role in the expression of age-dependent heritabilities for diameter growth, but not for height in Pinus radiata (Lin et al. 2013). Replicating our analysis in older trees would be interesting to further assess patterns of association between within-population genetic variation and environmental heterogeneity, and their underlying causes.

Finally, a last explanation is related to the different biological features between lodgepole pine and maritime pine. Lodgepole pine has extensive gene flow and low population structure (F_*ST*_ = 0.016 in Yeaman et al. 2016) while maritime pine shows restricted gene flow with strong population structure (at least six distinct gene pools and F_*ST*_ = 0.112; Jaramillo-Correa et al. 2015; our study) and fragmented distribution (Alberto et al. 2013). Pollen dispersal kernels in maritime pine are highly leptokurtic, as for other wind-pollinated pines (Schuster and Mitton 2000, Robledo-Arnuncio and Gil 2005), with estimated mean dispersal distances from 78.4 to 174.4m (de-Lucas et al. 2008). Interestingly, McDonald and Yeaman (2018) showed that high levels of quantitative genetic variance can be maintained when a trait is under stabilizing selection only at intermediate levels of migration. Migration rates in maritime pine may therefore not be strong enough to compensate for the purifying effect of natural selection in heterogeneous environments, especially in mountainous areas which may represent barriers to gene flow and where populations are more isolated (see González-Martínez et al. 2007 for maritime pine). Meanwhile, in the homogeneous plateaus of the Landes forest and central Spain, natural selection may be low because conditions are more favorable, and these populations are less isolated, which may maintain genetic variation at levels similar to those of populations in heterogeneous landscapes. Investigating local adaptation and gene flow at microgeographic scales in natural populations of maritime pine located in both homogeneous and heterogeneous environments would be highly valuable to understand why environmental heterogeneity does not seem to play a major role in maintaining genetic variation in this species. Moreover, conducting similar analyses in sister species such as Scots pine, with low population genetic structure and continuous populations (Alberto et al. 2013), could help to determine whether genetic variation in forest tree populations experiencing higher migration rates are more prone to be impacted by environmental heterogeneity.

### 4.3 Link to fitness and genetic constraints may explain the different patterns across traits

Height was the only trait that showed a consistent association between within-population genetic variation and climate harshness. This pattern supports the hypothesis that natural selection mainly depletes genetic variation of traits most directly related to fitness. Indeed, height can be seen as the end-product of multiple ecophysiological processes (Grattapaglia et al. 2009). Taller trees perform better in the competition for light, water and nutrients, and are therefore more likely to have higher fecundity (Rehfeldt et al. 1999, Wu and Ying 2004, Aitken and Bemmels 2015) and lower mortality (Wyckoff and Clark 2002, Zhu et al. 2017). However, taller trees are also more susceptible to spring and fall cold injury (Howe et al. 2003) and to drought (Bennett et al. 2015, McDowell and Allen 2015, Stovall et al. 2019). In maritime pine, effective reproductive success (i.e. the number of successfully established offspring) is related to tree size. Indeed, González-Martínez et al. (2006) found a significant positive female selection gradient for diameter (height was not tested, but diameter and height are strongly correlated in conifers; see, for example, Fig. 1 in Castedo-Dorado et al. 2005 for maritime pine) and suggested that offspring mothered by bigger trees could have a selective advantage due to better quality seeds favouring resilience in the face of severe summer droughts and microsite variation. This evidence also supports the idea of height as a relevant fitness component in maritime pine.

Although less directly related to fitness than height, leaf phenology-related traits exhibit steep adaptation gradients in forest trees and have a relatively high heritability, e.g. 0.15-0.51 for bud burst in pedunculate oak (Scotti-Saintagne et al. 2004), 0.45-1 in Sitka spruce (Alfaro et al. 2000) and 0.54 for bud burst and 0.30 for the duration of bud burst in our study in maritime pine. Gauzere et al. (2020) showed that both the mean and the variance of leaf phenology-related traits varied along an altitudinal gradient in natural oak populations, with populations at high altitude having a narrower fitness peak. We might therefore have expected lower genetic variation for leaf phenology-related traits in populations experiencing severe cold events (and at higher altitude), as found along an altitudinal gradient in sessile oak for bud phenology (Alberto et al. 2011). However, such association may be hidden in common gardens with different climates from those of the populations’ location, because of the release of high levels of cryptic genetic variation (Schlichting 2008). Moreover, phenology-related traits can show opposite genetic clines in common gardens and natural populations (e.g. Vitasse et al. 2009). Estimating genetic parameters of phenology-related traits directly in the field, which is now technically possible by using large genomic datasets and advanced statistical methodologies (Gienapp et al. 2017), may therefore be necessary to investigate potential associations between within-population genetic variation and climate harshness, or other selective pressures.

Importantly, theoretical work suggests that much of the genetic variation associated with a trait is likely maintained by pleiotropic effects, which are independent of the selection on that trait, implying that stabilizing selection can only act on a reduced number of independent dimensions in the trait space (Barton 1990, Walsh and Lynch 2018). As we used univariate models, we cannot exclude that the likely associations with height genetic variation originate from genetic correlation with other traits under selection, or that the lack of association with other traits (notably functional traits such as *δ*^13^C) does not originate from genetic constraints (Walsh and Blows 2009). For example, in maritime pine, trait canalisation and genetic constraints may explain low quantitative genetic differentiation for hydraulic traits (e.g. P50, the xylem pressure inducing 50% loss of hydraulic conductance; Lamy et al. 2014), and sapling height was found to be either positively or negatively associated with disease susceptibility depending on the pathogen (e.g. necrosis length caused by *Diplodia sapinea* or *Armillaria ostoyae*, respectively; Hurel et al. 2019). Trade-offs between traits may also explain the unexpected association between minimum temperatures and high genetic variation for SLA, as, for instance, SLA is known to be positively correlated with leaf life span, low assimilation rates and nutrient retention, i.e. traits linked to conservation of acquired resources (Ackerly et al. 2002).

## 5 Conclusion

Our manuscript contributes to the current debate on the maintenance of quantitative genetic variation within populations by providing empirical support for the role of natural selection in decreasing genetic variation. Indeed, our results consistently showed that genetic variation for height is lower in maritime pine populations experiencing severe cold events (i.e. experiencing stronger selection). Surprisingly, we found no association between environmental heterogeneity at the regional scale and within-population genetic variation for several traits; whether for technical reasons (e.g. sample size, spatial scale considered) or for genuine biological reasons (e.g. too low migration), it would be worth further exploration. Indeed, understanding the evolutionary forces shaping within-population genetic variation could shed light on how populations adapt to their local environment, thereby providing insight into how they may respond to future changes in environmental conditions.

## 6 Acknowledgments

We thank A. Saldaña, F. del Caño, E. Ballesteros and D. Barba (INIA) and the ‘Unitóe Expérimentale Forêt Pierroton’ (UEFP, INRAE; doi:10.15454/1.5483264699193726E12) for field assistance (plantation and measurements). Data used in this research are part of the Spanish Network of Genetic Trials (GENFORED, http://www.genfored.es). We thank all persons and institutions linked to the establishment and maintenance of field trials used in this study. We are very grateful to Ricardo Alía and Juan Majada who initiated and supervised the establishment of the CLONAPIN network and provided the progeny test height data that we used in the validation analysis. We thank Maurizio Marchi for kindly providing the raster files of the climate variables corresponding to the desired time period and spatial extent. JA was funded by the University of Bordeaux (ministerial grant). This study was funded by the ‘Initiative d’Excellence (IdEx) de l’Université de Bordeaux - Chaires d’installation 2015’ (EcoGenPin) and the European Union’s Horizon 2020 research and innovation programme under grant agreements No 773383 (B4EST) and No 862221 (FORGENIUS).

