## Supplementary Information for "Extreme climatic events but not environmental heterogeneity shape within-population genetic variation in maritime pine"

**1 The data** **2**

**2 Model equation and priors** **14**

**3 Model accuracy on simulated data** **15**

**4 $\beta_X$ interpretation** **15**

**5 Model outputs** **17**

**8 Changes since preregistration** **33**

### 1 The data

#### 1.1 Phenotypic data

##### 1.1.1 Details of the eight phenotypic traits

| Traits | Common gardens | Dates of measurement | Tree age | Survival | Units | Trees | Populations | Clones | Transf. |
|---|---|---|---|---|---|---|---|---|---|
| Height | Portugal | October 2012 | 20 | 0.66 | mm | 2746 | 33 | 521 | - |
| Height | Bordeaux | November 2013 | 25 | 0.97 | mm | 3238 | 33 | 430 | - |
| Height | Bordeaux | November 2018 | 85 | 0.96 | mm | 3209 | 33 | 430 | - |
| Height | Asturias | November 2012 | 21 | 0.96 | mm | 3973 | 33 | 522 | - |
| Mean bud burst date | Bordeaux | 2013, 2014, 2015, 2017 | - | - | °C-day | 3175 | 33 | 430 | center |
| Mean duration of bud burst | Bordeaux | 2014, 2015, 2017 | - | - | °C-day | 3187 | 33 | 430 | center |
| Specific Leaf Area | Portugal | - | - | - | $m^2/kg$ | 2642 | 33 | 520 | log |
| $\delta^{13}C$ | Portugal | - | - | - | ‰ | 1939 | 33 | 517 | center |

**Table S1:** Information about the phenotypic traits. Tree age is in months. Survival is the proportion of survival in the common gardens at the measurement date. Some variables were log-transformed (*log*) or mean-centered (*center*) prior to analyses, which is indicated in the *Transf.* column.

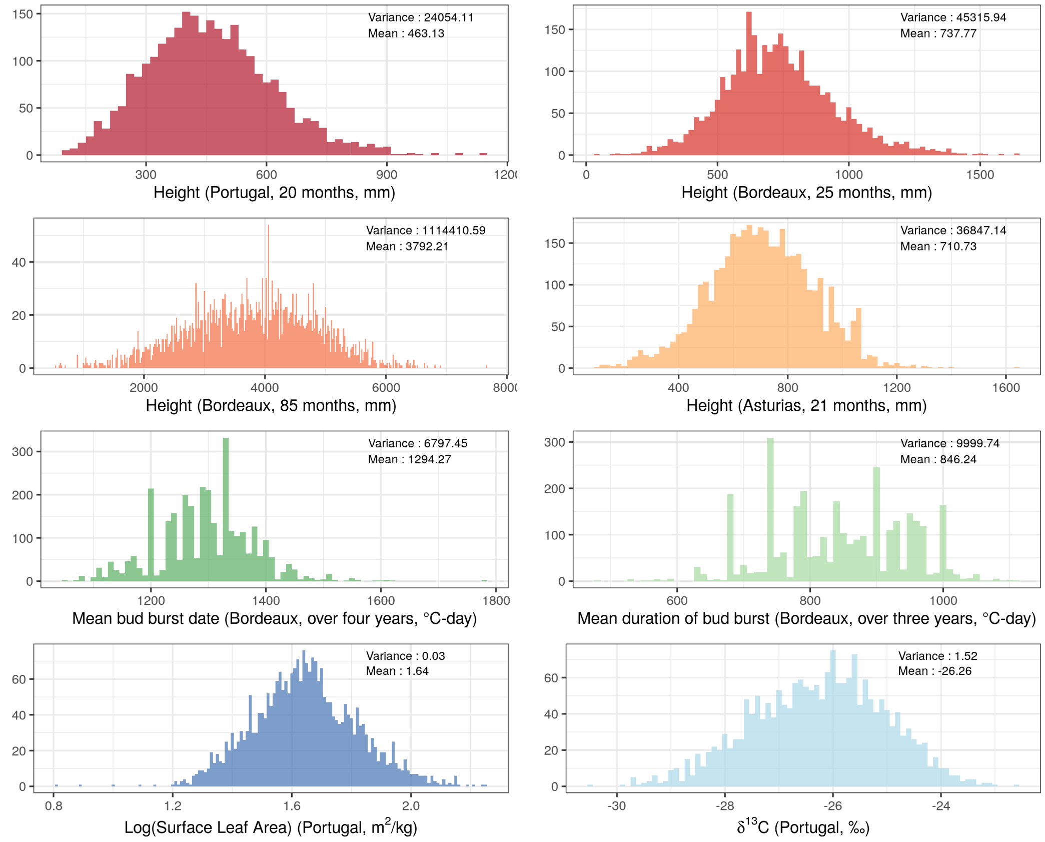

**Figure S1:** Distribution of the phenotypic traits used in the study.

##### 1.1.2 Population-specific distributions, means and variances

| **Pop.** | **Trait means** | | | | | | | | **Trait variances** | | | | | | | |
|---|---|---|---|---|---|---|---|---|---|---|---|---|---|---|---|---|
| | Ht (Portugal) | Ht (Bordeaux 2013) | Ht (Bordeaux 2018) | Ht (Asturias) | SLA | $\delta^{13}C$ | meanBB | meanDBB | Ht (Portugal) | Ht (Bordeaux 2013) | Ht (Bordeaux 2018) | Ht (Asturias) | SLA | $\delta^{13}C$ | meanBB | meanDBB |
| ALT | 525.56 | 760.19 | 4275.38 | 764.43 | 5.08 | -25.94 | 1333.05 | 883.19 | 25044.03 | 25994.08 | 518190.0 | 43958.36 | 0.56 | 1.26 | 10179.89 | 11621.08 |
| ARM | 505.48 | 793.33 | 4450.16 | 818.89 | 5.19 | -26.01 | 1282.95 | 859.95 | 14693.67 | 30154.84 | 416420.9 | 27155.20 | 0.53 | 0.75 | 5787.12 | 10512.04 |
| ARN | 423.44 | 651.87 | 3302.00 | 652.35 | 5.19 | -26.83 | 1309.37 | 861.13 | 20251.08 | 26534.21 | 487704.6 | 21763.14 | 0.96 | 1.68 | 1199.55 | 5285.84 |
| BAY | 394.29 | 618.36 | 3054.03 | 592.79 | 5.18 | -26.49 | 1299.02 | 829.97 | 15606.98 | 24433.97 | 461502.5 | 20475.10 | 0.96 | 1.91 | 5347.63 | 7758.00 |
| BON | 421.33 | 600.43 | 3273.83 | 702.61 | 5.37 | -26.84 | 1197.00 | 752.85 | 15994.80 | 30225.90 | 1159572.0 | 26013.68 | 1.08 | 1.12 | 3843.38 | 7596.92 |
| CAD | 481.58 | 847.76 | 4631.79 | 781.60 | 5.34 | -25.79 | 1342.52 | 898.14 | 32160.59 | 55641.88 | 894272.5 | 42048.76 | 0.85 | 1.11 | 13379.00 | 8728.85 |
| CAR | 405.52 | 505.32 | 2321.30 | 584.58 | 4.81 | -26.74 | 1282.36 | 813.30 | 18268.47 | 18912.40 | 396287.2 | 11029.61 | 0.58 | 1.22 | 9461.82 | 8783.43 |
| CAS | 501.67 | 753.01 | 4202.78 | 769.32 | 5.07 | -25.90 | 1310.38 | 877.46 | 28617.67 | 35004.68 | 554102.0 | 30359.25 | 1.00 | 1.01 | 6072.29 | 12050.36 |
| CEN | 484.82 | 663.51 | 3348.65 | 723.14 | 4.91 | -26.32 | 1272.32 | 849.04 | 25174.51 | 31090.09 | 648606.5 | 26016.07 | 0.58 | 1.36 | 7633.28 | 15793.70 |
| COC | 406.47 | 587.13 | 2735.71 | 588.77 | 5.22 | -26.47 | 1320.11 | 848.06 | 17647.00 | 25181.40 | 573367.0 | 25089.71 | 0.89 | 1.39 | 6714.06 | 7676.71 |
| COM | 516.92 | 809.33 | 3220.71 | 756.13 | 4.86 | -26.14 | 1260.71 | 836.87 | 22918.15 | 35020.23 | 342303.2 | 57311.18 | 0.53 | 0.83 | 4511.64 | 11303.29 |
| CUE | 402.29 | 587.38 | 2793.39 | 582.10 | 5.06 | -26.35 | 1313.72 | 851.46 | 15749.06 | 29948.47 | 745261.0 | 19274.90 | 1.02 | 1.76 | 7290.54 | 8915.74 |
| HOU | 549.58 | 841.95 | 4578.18 | 831.13 | 5.32 | -25.79 | 1287.73 | 832.88 | 30575.88 | 36168.73 | 683021.5 | 37114.18 | 0.80 | 1.09 | 3998.32 | 8832.28 |
| LAM | 443.56 | 706.79 | 3824.82 | 736.38 | 5.37 | -26.25 | 1322.51 | 893.70 | 18614.34 | 28385.84 | 503323.6 | 35455.80 | 0.90 | 1.44 | 4263.94 | 6633.86 |
| LEI | 467.30 | 765.64 | 4156.02 | 743.20 | 5.12 | -26.34 | 1317.84 | 880.28 | 23362.76 | 44161.14 | 799974.2 | 41372.76 | 0.81 | 1.29 | 8737.24 | 11448.55 |
| MIM | 447.76 | 783.87 | 4182.64 | 718.20 | 5.34 | -26.05 | 1283.85 | 832.76 | 22395.23 | 51903.59 | 755436.3 | 33853.99 | 0.91 | 1.94 | 6008.16 | 8888.56 |
| OLB | 445.60 | 700.35 | 3410.89 | 712.97 | 5.24 | -26.43 | 1285.80 | 817.56 | 15336.79 | 23496.30 | 487763.2 | 24782.38 | 0.92 | 1.28 | 5575.52 | 9353.93 |
| OLO | 549.10 | 803.57 | 4390.16 | 828.55 | 5.41 | -25.63 | 1287.50 | 829.00 | 22867.78 | 47499.69 | 534286.6 | 29013.66 | 0.84 | 1.08 | 4316.43 | 9240.09 |
| ORI | 375.58 | 667.40 | 2935.64 | 639.17 | 5.40 | -26.92 | 1239.57 | 797.85 | 15107.03 | 21736.00 | 391840.3 | 25020.48 | 1.10 | 0.94 | 4685.37 | 7988.44 |
| PET | 525.13 | 829.51 | 4471.39 | 781.10 | 5.29 | -25.81 | 1306.26 | 864.40 | 24369.26 | 38187.87 | 665897.4 | 41792.11 | 0.67 | 1.53 | 3723.47 | 10308.61 |
| PIA | 494.60 | 838.62 | 4163.58 | 783.88 | 5.22 | -26.51 | 1201.04 | 793.13 | 28122.80 | 44180.50 | 558897.3 | 35430.62 | 0.84 | 1.39 | 3697.91 | 9221.80 |
| PIE | 435.18 | 630.77 | 3318.63 | 620.29 | 5.43 | -26.26 | 1278.11 | 847.36 | 19269.41 | 23336.65 | 472940.1 | 30191.09 | 1.02 | 1.37 | 3240.16 | 7546.22 |
| PLE | 468.99 | 770.15 | 4044.84 | 683.62 | 5.42 | -25.92 | 1306.42 | 870.85 | 25399.24 | 46364.32 | 868825.2 | 33619.27 | 1.12 | 1.23 | 5312.36 | 7901.84 |
| PUE | 520.26 | 865.10 | 4460.59 | 775.65 | 4.98 | -26.10 | 1353.32 | 902.72 | 33564.79 | 45797.49 | 841781.7 | 37674.17 | 0.72 | 1.11 | 5397.23 | 8249.97 |
| QUA | 458.63 | 706.61 | 3265.79 | 689.08 | 4.92 | -26.74 | 1304.61 | 862.73 | 20764.43 | 25415.59 | 679962.7 | 27766.85 | 0.61 | 1.53 | 5845.11 | 8570.03 |
| SAC | 436.84 | 779.80 | 4317.55 | 699.71 | 5.17 | -25.69 | 1333.98 | 882.42 | 10276.24 | 26602.00 | 665431.4 | 40020.50 | 0.56 | 0.57 | 10387.92 | 11888.99 |
| SAL | 396.90 | 582.90 | 2915.65 | 592.48 | 5.33 | -26.74 | 1253.29 | 802.65 | 20116.52 | 15944.42 | 339030.8 | 19588.04 | 0.95 | 1.01 | 5040.26 | 6804.81 |
| SEG | 453.87 | 819.44 | 4484.53 | 728.92 | 5.12 | -26.06 | 1320.39 | 878.22 | 19458.77 | 47514.06 | 767773.7 | 26366.13 | 0.81 | 1.07 | 6328.79 | 10553.06 |
| SIE | 445.13 | 860.65 | 4266.96 | 755.56 | 5.42 | -26.11 | 1337.77 | 887.32 | 28151.96 | 49975.12 | 431452.8 | 29270.25 | 0.88 | 1.34 | 5703.93 | 7581.65 |
| STJ | 535.06 | 893.84 | 4559.44 | 794.76 | 5.43 | -25.92 | 1296.49 | 838.08 | 23677.59 | 45913.57 | 632962.2 | 37600.18 | 0.81 | 1.60 | 6258.52 | 10697.50 |
| TAM | 339.09 | 570.29 | 2250.45 | 555.13 | 5.76 | -27.82 | 1261.34 | 813.01 | 12214.55 | 16014.62 | 365171.0 | 23907.35 | 0.79 | 0.89 | 3427.87 | 7982.86 |
| VAL | 431.30 | 672.13 | 3393.73 | 656.09 | 5.13 | -26.56 | 1258.81 | 825.76 | 16849.74 | 43523.72 | 860272.1 | 23268.04 | 0.92 | 1.06 | 6513.53 | 10003.81 |
| VER | 503.62 | 813.65 | 4401.61 | 780.73 | 5.46 | -25.75 | 1325.48 | 864.43 | 22141.52 | 44992.37 | 680372.1 | 34422.39 | 0.73 | 1.57 | 4422.45 | 8800.61 |

**Table S2:** Population-specific means and variances of the eight phenotypic traits (from left to right): height in Portugal (October 2012), height in Bordeaux (France, November 2013), height in Bordeaux (France, November 2018), height in Asturias (Spain, November 2012), mean bud burst date in Bordeaux (over the years 2013, 2014, 2015 and 2017), mean duration of bud burst in Bordeaux (over the years 2014, 2015 and 2017), specific leaf area in Portugal and $\delta^{13}C$ in Portugal.

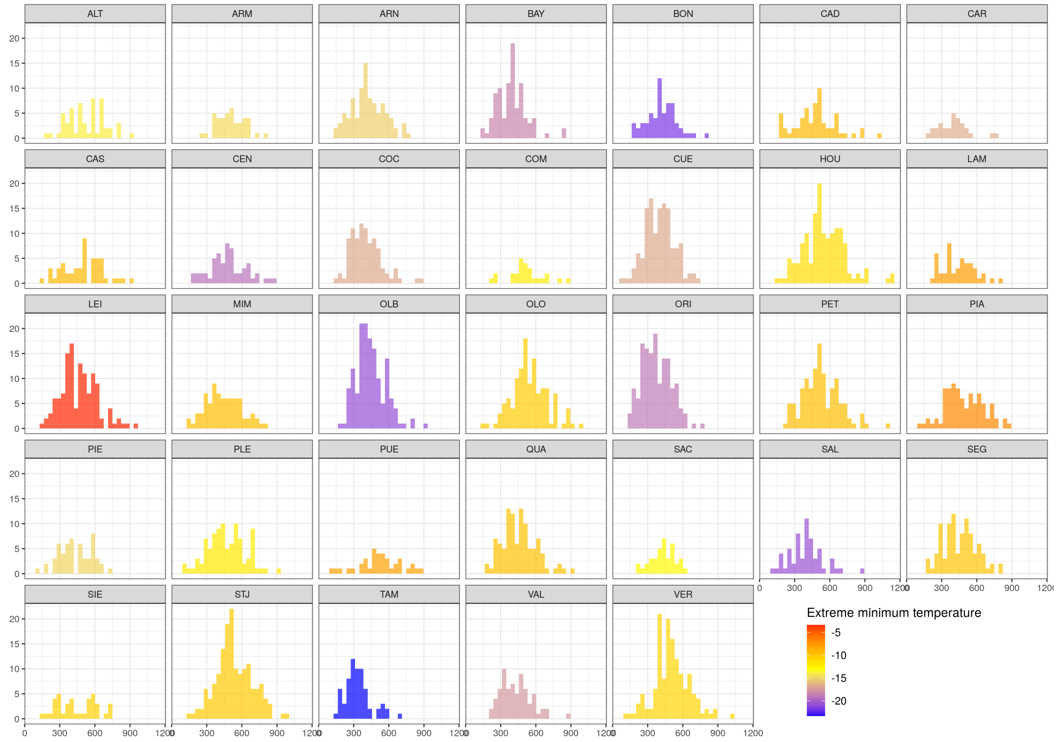

**Figure S2:** Population-specific height distributions in Portugal (October 2012). The color gradient corresponds to the extreme minimum temperature in the population location over the period 1901-1950.

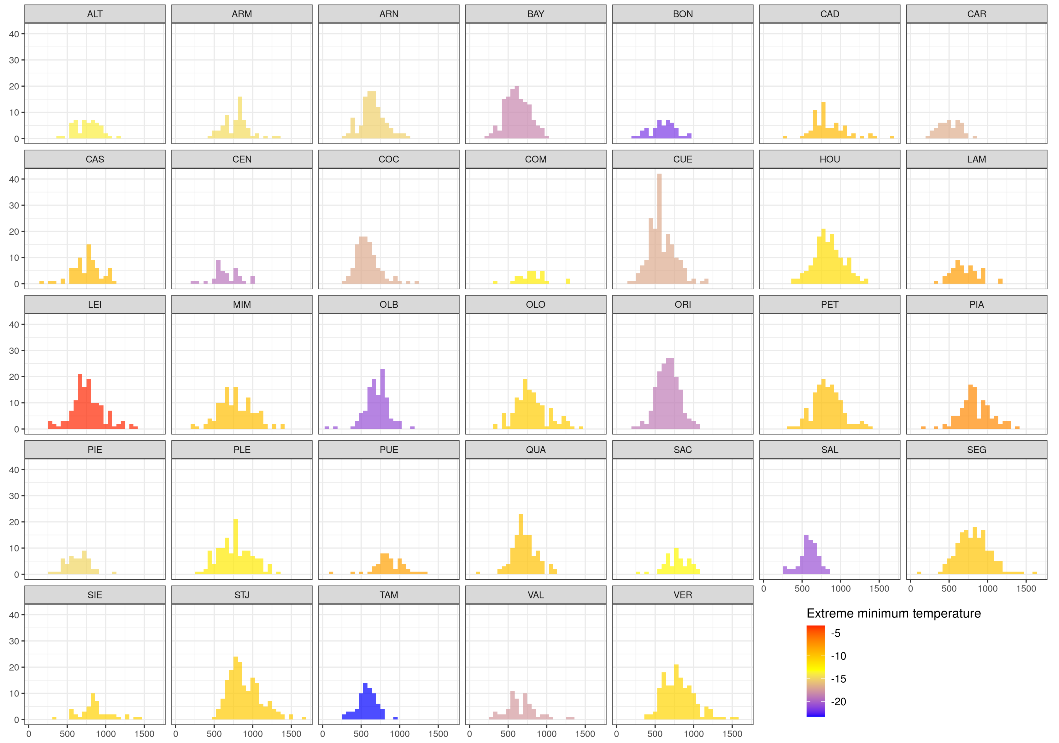

**Figure S3:** Population-specific height distributions in Bordeaux (France, November 2013). The color gradient corresponds to the extreme minimum temperature in the population location over the period 1901-1950.

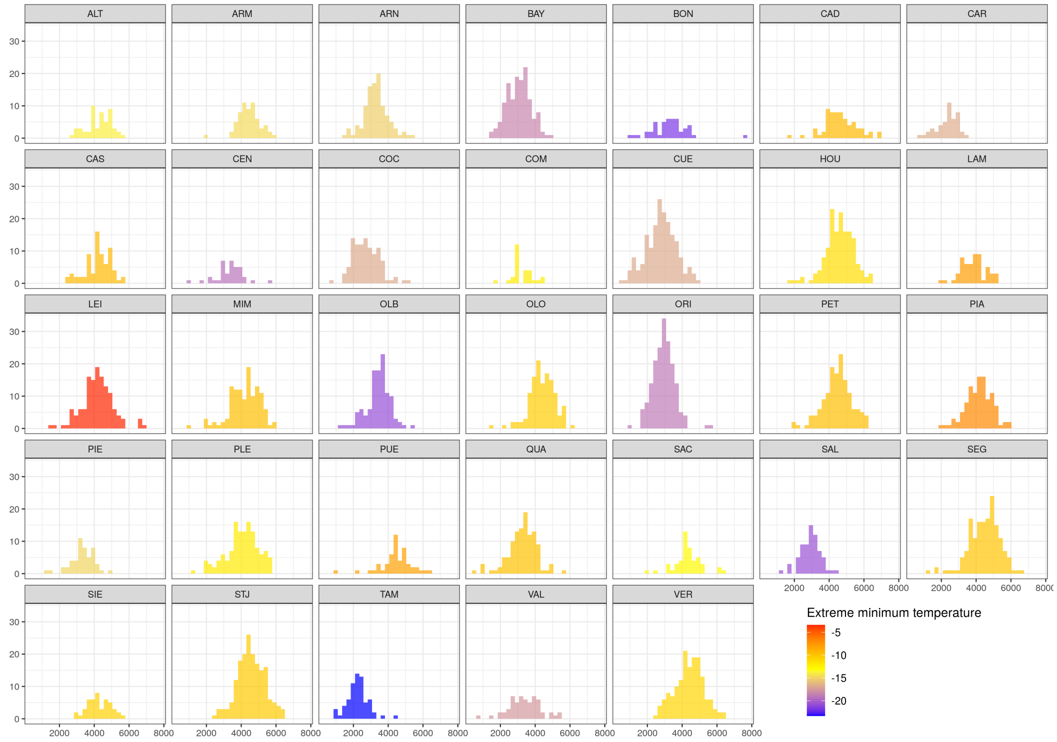

**Figure S4:** Population-specific height distributions in Bordeaux (France, November 2018). The color gradient corresponds to the extreme minimum temperature in the population location over the period 1901-1950.

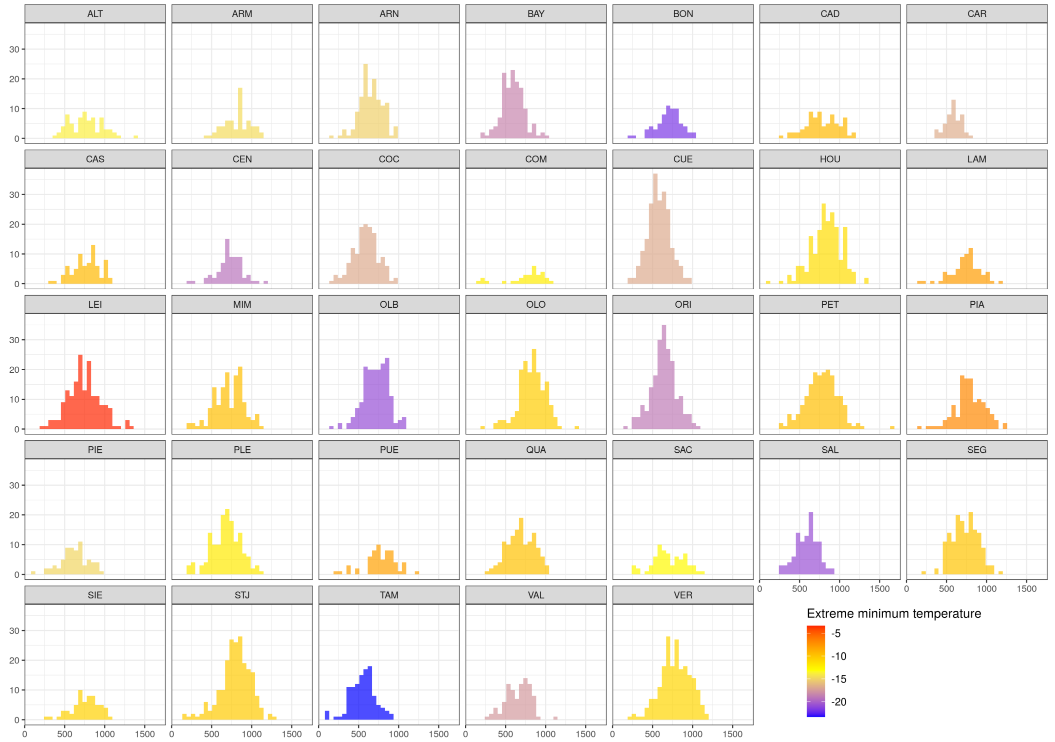

**Figure S5:** Population-specific height distributions in Asturias (Spain, November 2012). The color gradient corresponds to the extreme minimum temperature in the population location over the period 1901-1950.

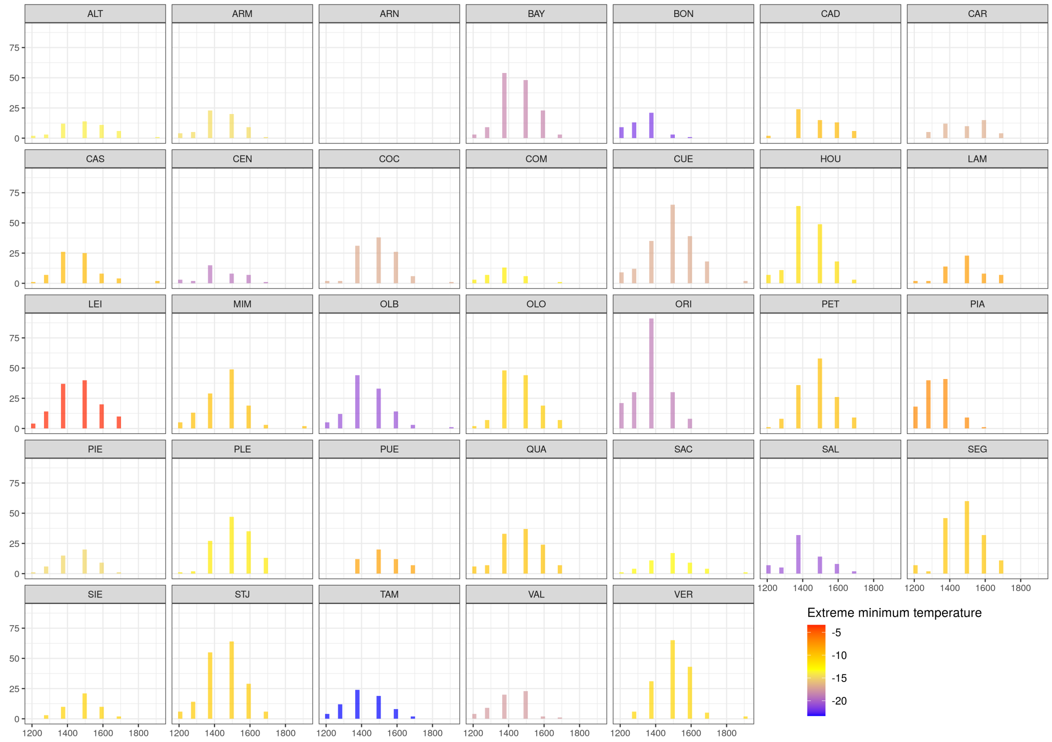

**Figure S6:** Population-specific distributions of the mean bud burst date in Bordeaux (France) averaged over the years 2013, 2014, 2015 and 2017. The color gradient corresponds to the extreme minimum temperature in the population location over the period 1901-1950.

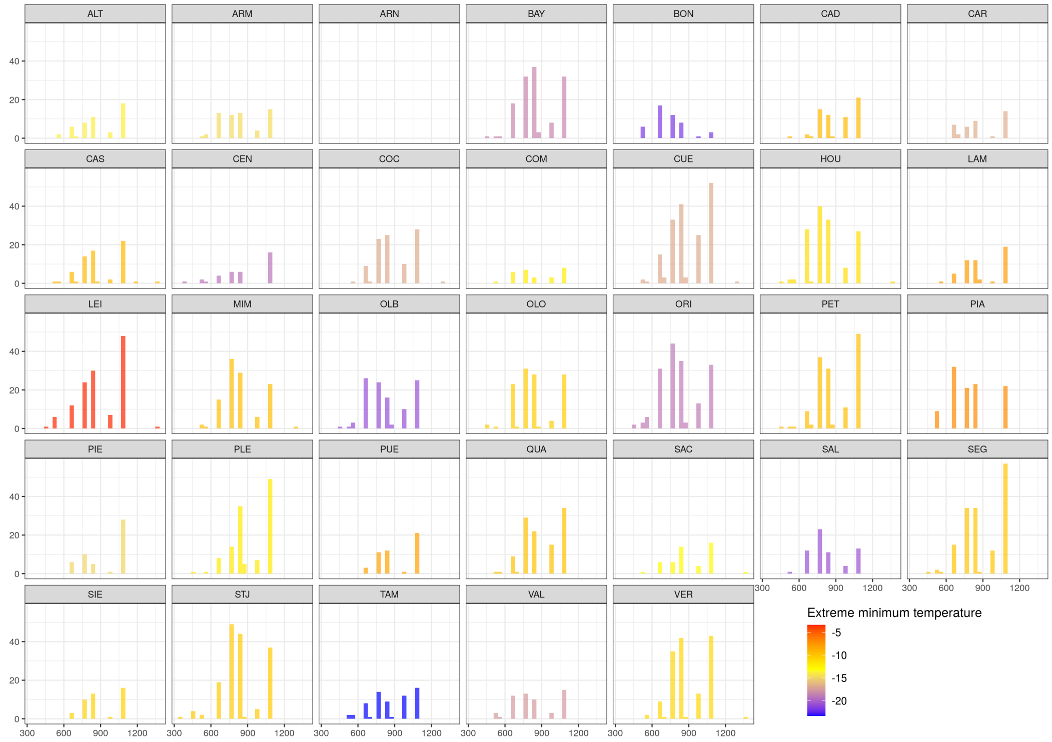

**Figure S7:** Population-specific distributions of the mean duration of the bud burst date in Pieroton (France) averaged over the years 2014, 2015 and 2017. The color gradient corresponds to the extreme minimum temperature in the population location over the period 1901-1950.

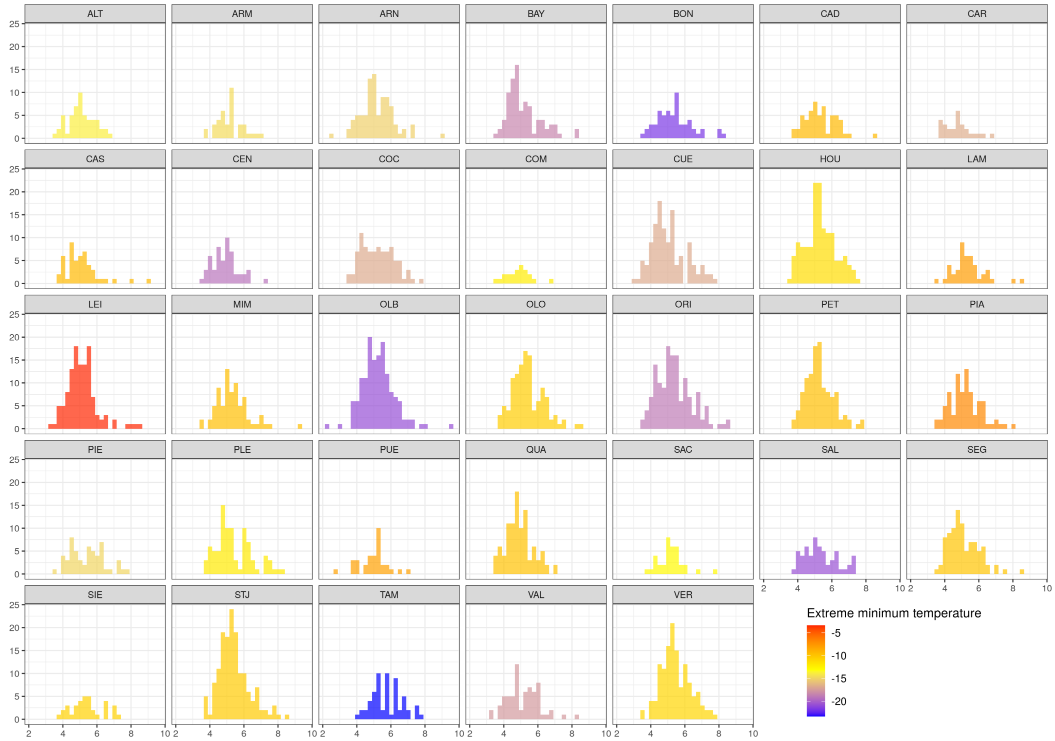

**Figure S8:** Population-specific distributions of the specific leaf area (SLA) in Portugal. The color gradient corresponds to the extreme minimum temperature in the population location over the period 1901-1950.

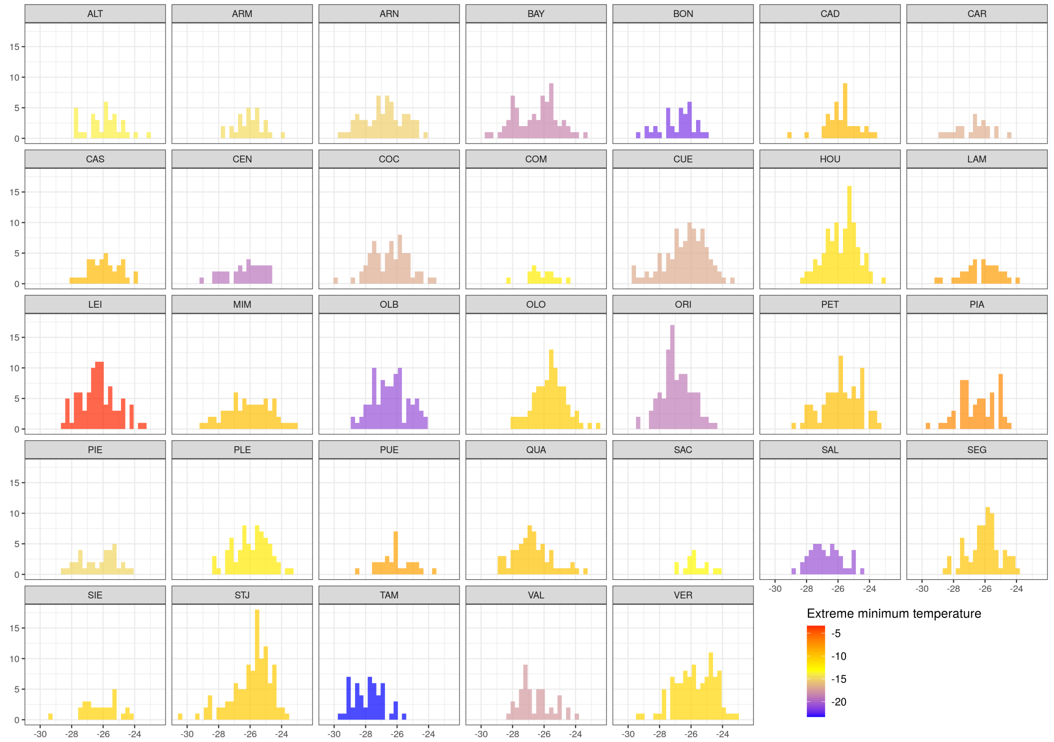

**Figure S9:** Population-specific distributions of the isotope discrimination ($\delta^{13}C$) in Portugal. The color gradient corresponds to the extreme minimum temperature in the population location over the period 1901-1950.

#### 1.2 Environment of the common gardens

| Coordinates, soil and topographic variables | Units | Asturias | Bordeaux | Portugal |
|---|---|---|---|---|
| Latitude | degrees | 43.42 | 44.75 | 40.11 |
| Longitude | degrees | -6.54 | -0.78 | -7.48 |
| Topographic ruggedness index | unitless | 17.18 | 0.81 | 19.76 |
| Depth available to roots | cm | 30.00 | 70.00 | 70.00 |
| Clay content in the topsoil (0-30 cm) | % | 20.00 | 4.00 | 10.00 |
| Sand content in the topsoil (0-30 cm) | % | 49.00 | 87.00 | 75.00 |
| Silt content in the topsoil (0-30 cm) | % | 31.00 | 9.00 | 15.00 |

**Table S3:** Values of the geographical coordinates, soil and topographic variables of the three common gardens.

| Annual climatic variables | Units | Asturias | Bordeaux | Portugal |
|---|---|---|---|---|
| Mean Coldest Month Temperature (MCMT) | °C | 5.80 | 5.60 | 4.70 |
| Mean Warmest Month Temperature (MWMT) | °C | 17.75 | 19.70 | 19.75 |
| Temperature difference (MWMT-MCMT) | °C | 11.95 | 14.10 | 15.05 |
| Extreme minimum temperature (**EMT**) | °C | -12.50 | -13.30 | -14.30 |
| Mean summer precipitation | mm | 423.00 | 496.00 | 370.00 |
| Mean spring precipitation | mm | 242.00 | 226.00 | 271.00 |
| Summer heat moisture index (**SHM**) | °C/mm | 134.47 | 105.91 | 274.31 |

**Table S4:** Values of the annual climatic variables in the three common gardens.

| Monthly climatic variables | Units | Asturias | Bordeaux | Portugal |
|---|---|---|---|---|
| Minimum temperature - January | °C | 2.50 | 2.00 | 1.50 |
| Minimum temperature - February | °C | 3.00 | 2.50 | 1.60 |
| Minimum temperature - March | °C | 4.20 | 3.80 | 3.10 |
| Minimum temperature - April | °C | 5.40 | 6.30 | 4.70 |
| Minimum temperature - May | °C | 7.70 | 9.20 | 7.30 |
| Minimum temperature - June | °C | 10.50 | 12.10 | 10.90 |
| Minimum temperature - July | °C | 12.30 | 13.90 | 13.10 |
| Minimum temperature - August | °C | 12.70 | 13.80 | 13.40 |
| Minimum temperature - September | °C | 11.30 | 11.80 | 11.30 |
| Minimum temperature - October | °C | 8.30 | 8.60 | 8.20 |
| Minimum temperature - November | °C | 5.40 | 4.80 | 4.20 |
| Minimum temperature - December | °C | 3.50 | 2.80 | 2.20 |
| Maximum temperature - January | °C | 9.10 | 9.20 | 7.90 |
| Maximum temperature - February | °C | 10.40 | 11.10 | 8.80 |
| Maximum temperature - March | °C | 12.30 | 13.90 | 11.10 |
| Maximum temperature - April | °C | 14.10 | 16.60 | 13.90 |
| Maximum temperature - May | °C | 16.70 | 20.00 | 17.20 |
| Maximum temperature - June | °C | 20.20 | 23.30 | 22.10 |
| Maximum temperature - July | °C | 22.50 | 25.50 | 25.70 |
| Maximum temperature - August | °C | 22.80 | 25.60 | 26.10 |
| Maximum temperature - September | °C | 20.90 | 23.30 | 22.00 |
| Maximum temperature - October | °C | 16.70 | 18.30 | 16.60 |
| Maximum temperature - November | °C | 12.20 | 12.80 | 11.20 |
| Maximum temperature - December | °C | 9.70 | 9.50 | 8.40 |
| Total precipitation - January | mm | 88.00 | 83.00 | 112.00 |
| Total precipitation - February | mm | 79.00 | 74.00 | 122.00 |
| Total precipitation - March | mm | 93.00 | 76.00 | 120.00 |
| Total precipitation - April | mm | 72.00 | 70.00 | 78.00 |
| Total precipitation - May | mm | 77.00 | 80.00 | 73.00 |
| Total precipitation - June | mm | 50.00 | 63.00 | 45.00 |
| Total precipitation - July | mm | 34.00 | 50.00 | 11.00 |
| Total precipitation - August | mm | 37.00 | 54.00 | 9.00 |
| Total precipitation - September | mm | 61.00 | 82.00 | 52.00 |
| Total precipitation - October | mm | 92.00 | 97.00 | 102.00 |
| Total precipitation - November | mm | 100.00 | 98.00 | 122.00 |
| Total precipitation - December | mm | 113.00 | 103.00 | 124.00 |
| Hargreaves climatic moisture deficit - January | mm | 17.68 | 17.88 | 18.17 |
| Hargreaves climatic moisture deficit - February | mm | 26.56 | 29.31 | 25.17 |
| Hargreaves climatic moisture deficit - March | mm | 46.40 | 57.06 | 44.75 |
| Hargreaves climatic moisture deficit - April | mm | 68.38 | 82.97 | 72.07 |
| Hargreaves climatic moisture deficit - May | mm | 92.24 | 115.64 | 101.41 |
| Hargreaves climatic moisture deficit - June | mm | 115.38 | 137.41 | 134.79 |
| Hargreaves climatic moisture deficit - July | mm | 130.36 | 151.65 | 166.13 |
| Hargreaves climatic moisture deficit - August | mm | 115.21 | 134.58 | 151.93 |
| Hargreaves climatic moisture deficit - September | mm | 79.08 | 92.08 | 92.73 |
| Hargreaves climatic moisture deficit - October | mm | 45.16 | 50.68 | 48.35 |
| Hargreaves climatic moisture deficit - November | mm | 21.58 | 23.63 | 23.31 |
| Hargreaves climatic moisture deficit - December | mm | 14.70 | 14.68 | 16.03 |

**Table S5:** Values of the monthly climatic variables in the three common gardens.

#### 1.3 Potential drivers of the within-population genetic variation

##### 1.3.1 Population admixtures scores

| Population | Longitude | Latitude | gpNA | gpC | gpCS | gpFA | gpIA | gpSES | mainGP | A | D | $D_{fst}$ |
|---|---|---|---|---|---|---|---|---|---|---|---|---|
| CEN | -4.491 | 40.278 | 0.012 | 0.002 | 0.884 | 0.003 | 0.048 | 0.051 | gpCS | 0.116 | 0.031 | 0.011 |
| ARN | -5.116 | 40.195 | 0.010 | 0.002 | 0.955 | 0.008 | 0.010 | 0.014 | gpCS | 0.045 | 0.014 | 0.005 |
| ALT | -6.494 | 43.283 | 0.003 | 0.000 | 0.109 | 0.092 | 0.793 | 0.002 | gpIA | 0.207 | 0.061 | 0.019 |
| SAL | -3.063 | 41.835 | 0.010 | 0.004 | 0.944 | 0.027 | 0.008 | 0.007 | gpCS | 0.056 | 0.016 | 0.006 |
| COM | -3.954 | 36.834 | 0.240 | 0.028 | 0.126 | 0.011 | 0.039 | 0.557 | gpSES | 0.443 | 0.218 | 0.061 |
| CAD | -6.418 | 43.540 | 0.002 | 0.001 | 0.050 | 0.010 | 0.935 | 0.002 | gpIA | 0.065 | 0.018 | 0.005 |
| VAL | -4.311 | 40.516 | 0.012 | 0.003 | 0.940 | 0.007 | 0.013 | 0.025 | gpCS | 0.060 | 0.018 | 0.007 |
| MIM | -1.303 | 44.134 | 0.004 | 0.002 | 0.025 | 0.951 | 0.012 | 0.006 | gpFA | 0.049 | 0.014 | 0.006 |
| LEI | -8.957 | 39.783 | 0.004 | 0.003 | 0.511 | 0.007 | 0.473 | 0.002 | gpCS | 0.489 | 0.125 | 0.031 |
| BAY | -2.877 | 41.523 | 0.003 | 0.004 | 0.967 | 0.013 | 0.009 | 0.004 | gpCS | 0.033 | 0.009 | 0.003 |
| SIE | -6.493 | 43.528 | 0.003 | 0.001 | 0.076 | 0.015 | 0.903 | 0.001 | gpIA | 0.097 | 0.028 | 0.008 |
| LAM | -6.219 | 43.559 | 0.002 | 0.001 | 0.004 | 0.051 | 0.942 | 0.000 | gpIA | 0.058 | 0.020 | 0.007 |
| TAM | -5.017 | 33.600 | 0.932 | 0.000 | 0.027 | 0.000 | 0.039 | 0.001 | gpNA | 0.068 | 0.045 | 0.020 |
| COC | -4.498 | 41.255 | 0.017 | 0.005 | 0.831 | 0.060 | 0.044 | 0.044 | gpCS | 0.169 | 0.044 | 0.015 |
| OLO | -1.831 | 46.566 | 0.003 | 0.001 | 0.007 | 0.979 | 0.006 | 0.003 | gpFA | 0.021 | 0.007 | 0.003 |
| STJ | -2.029 | 46.764 | 0.003 | 0.002 | 0.028 | 0.946 | 0.017 | 0.004 | gpFA | 0.054 | 0.015 | 0.006 |
| CUE | -4.484 | 41.375 | 0.003 | 0.001 | 0.872 | 0.063 | 0.058 | 0.002 | gpCS | 0.128 | 0.030 | 0.009 |
| PET | -1.300 | 44.064 | 0.003 | 0.001 | 0.021 | 0.966 | 0.004 | 0.004 | gpFA | 0.034 | 0.009 | 0.004 |
| ORI | -2.351 | 37.531 | 0.248 | 0.005 | 0.028 | 0.001 | 0.010 | 0.708 | gpSES | 0.292 | 0.180 | 0.044 |
| SEG | -8.450 | 42.817 | 0.003 | 0.001 | 0.149 | 0.013 | 0.831 | 0.003 | gpIA | 0.169 | 0.046 | 0.012 |
| OLB | -0.623 | 40.173 | 0.080 | 0.009 | 0.777 | 0.006 | 0.003 | 0.125 | gpCS | 0.223 | 0.078 | 0.032 |
| QUA | -0.359 | 38.972 | 0.092 | 0.006 | 0.499 | 0.005 | 0.015 | 0.382 | gpCS | 0.501 | 0.142 | 0.056 |
| CAS | -6.983 | 43.501 | 0.001 | 0.000 | 0.005 | 0.001 | 0.992 | 0.001 | gpIA | 0.008 | 0.003 | 0.001 |
| PLE | -2.344 | 47.781 | 0.005 | 0.001 | 0.064 | 0.920 | 0.005 | 0.005 | gpFA | 0.080 | 0.020 | 0.008 |
| BON | -1.661 | 39.986 | 0.164 | 0.010 | 0.640 | 0.003 | 0.002 | 0.181 | gpCS | 0.360 | 0.139 | 0.057 |
| HOU | -1.150 | 45.183 | 0.004 | 0.001 | 0.026 | 0.960 | 0.007 | 0.002 | gpFA | 0.040 | 0.011 | 0.004 |
| VER | -1.091 | 45.552 | 0.003 | 0.002 | 0.018 | 0.972 | 0.004 | 0.001 | gpFA | 0.028 | 0.008 | 0.003 |
| ARM | -6.458 | 43.305 | 0.005 | 0.007 | 0.022 | 0.006 | 0.959 | 0.001 | gpIA | 0.041 | 0.015 | 0.005 |
| CAR | -4.277 | 41.172 | 0.001 | 0.001 | 0.904 | 0.060 | 0.023 | 0.011 | gpCS | 0.096 | 0.021 | 0.007 |
| PIE | 9.038 | 41.973 | 0.007 | 0.969 | 0.023 | 0.000 | 0.001 | 0.001 | gpC | 0.031 | 0.014 | 0.005 |
| PUE | -6.631 | 43.548 | 0.002 | 0.000 | 0.021 | 0.001 | 0.973 | 0.002 | gpIA | 0.027 | 0.008 | 0.003 |
| PIA | 9.465 | 42.021 | 0.004 | 0.974 | 0.010 | 0.001 | 0.008 | 0.003 | gpC | 0.026 | 0.012 | 0.005 |
| SAC | -8.364 | 42.118 | 0.004 | 0.001 | 0.291 | 0.003 | 0.699 | 0.002 | gpIA | 0.301 | 0.079 | 0.020 |

**Table S6:** Population admixture scores for the 33 provenances. The columns gpNA, gpC, gpCS, gpFA, gpIA and gpSES contain the proportion of belonging to the gene pools of Northern Africa, Corsica, Central Spain, French Atlantic region, Iberian Atlantic region and south-eastern Spain, respectively. A and D are the two population admixture scores used in the study. $D_{fst}$ is similar to D, as it was calculated by weighting the proportion of belonging from foreign gene pools by the pairwise $F_{ST}$ between the main and foreign gene pools, while D was calculated by weighting the proportion of belonging from foreign gene pools by the sum of the allele frequency divergence of the main and foreign gene pool from the common ancestral one (obtained from Jaramillo-Correa et al. 2015).

##### 1.3.2 Climate harshness and environmental heterogeneity indexes

| Environmental variables | Units |
|---|---|
| Monthly minimum temperature (12 variables) | °C |
| Monthly maximum temperature (12 variables) | °C |
| Monthly total precipitation (12 variables) | mm |
| Monthly Hargreaves climatic moisture deficit (12 variables) | mm |
| Mean Warmest Month Temperature (MWMT) | °C |
| Mean Coldest Month Temperature (MCMT) | °C |
| Temperature difference (MWMT-MCMT) | °C |
| Mean summer precipitation | mm |
| Mean spring precipitation | mm |
| Extreme minimum temperature (**EMT**) | °C |
| Summer heat moisture index (**SHM**) | °C/mm |
| Topographic ruggedness index | unitless |
| Depth available to roots | cm |
| Clay content in the topsoil (0-30 cm) | % |
| Sand content in the topsoil (0-30 cm) | % |
| Silt content in the topsoil (0-30 cm) | % |

**Table S7:** Climatic, topographic and soil variables used to describe the environmental heterogeneity around the population location (all variables) and the climate harshness at the population location (SHM and EMT). The climatic variables were averaged over the period 1901-1950, except the extreme minimum temperature (EMT) which corresponds to the minimum temperature over the same period.

**Forested areas:** In Europe, the forested areas were extracted from Copernicus ("Forest Type 2015" at 100-m resolution), which describes the land cover in 2015. We kept raster cells attributed to either broadleaved forests (1), coniferous forests (2) or mixed forests (3). The other raster cells were considered as non-forested areas. In Morocco, the forested areas were extracted from the GeoNetwork: Land cover of Morocco - Globcover Regional, which describes the land cover in 2005. Raster cells with the following LCCCode were considered as forested areas:

- 0003 / 0004 (Mosaic cropland vegetation)
- 0004 // 0003 (Mosaic vegetation / cropland)
- 21446 // 21450-121340 / 21454 (Mosaic forest or shrubland/grassland)
- 21450 (Closed to open shrubland)
- 21454 // 21446 // 21450 (Mosaic grassland/forest or shrubland)
- 21496 // 21497-15048 (Closed to open broadleaved evergreen or semidecidous forest)
- 21496-121340 // 21497-129401 (Closed (>40%) broadleaved evergreen or semidecidous forest)
- 21497-121340 (Closed (>40%) broadleaved decidous forest)
- 21497-15045 (Closed to open (>15%) mixed broadleaved decidous and needleaved evergreen forest)
- 21499-121340 (Closed (>40%) needeleaved evergreen forest)

- 21518 (Closed to open broadleaved decidous shrubland)

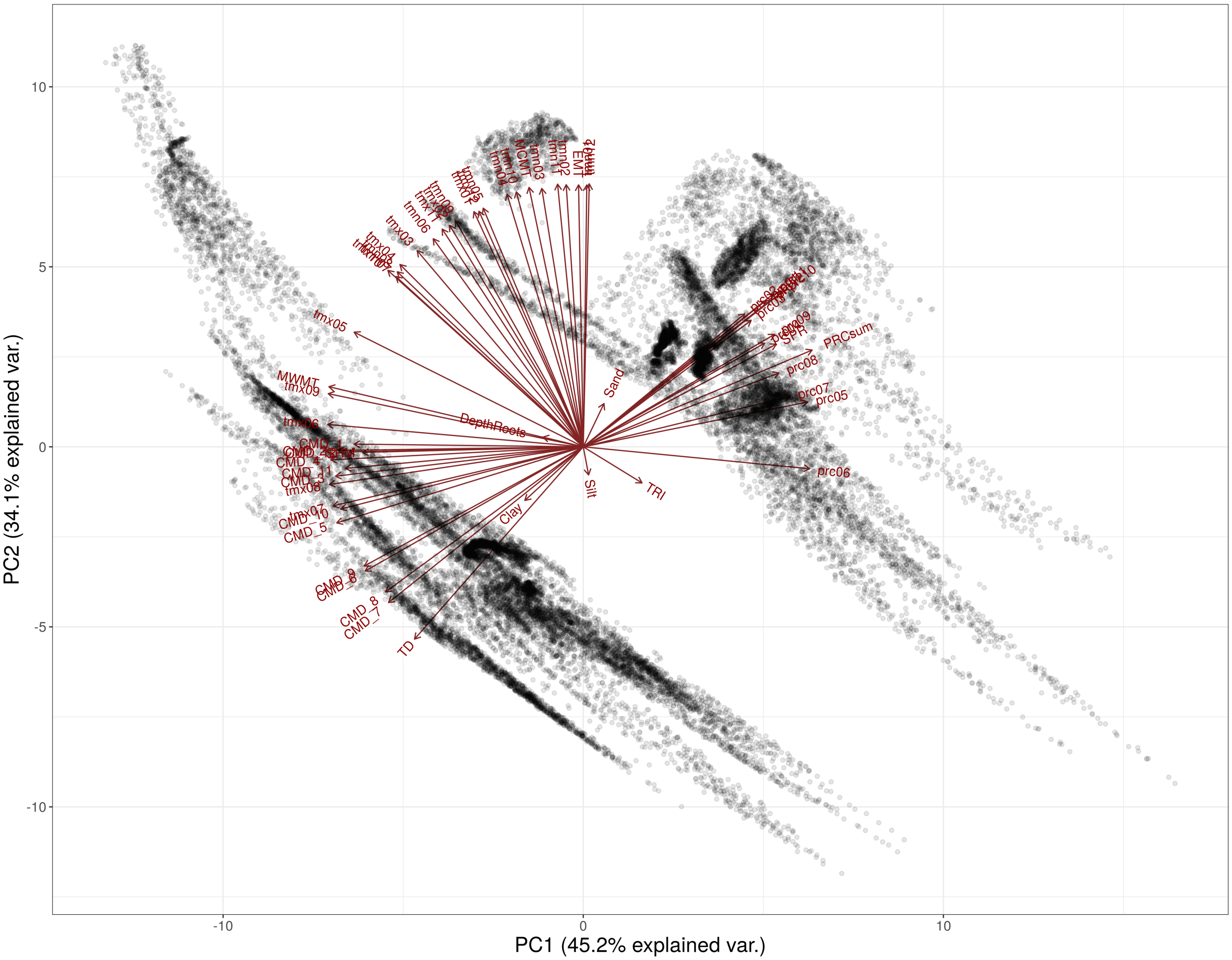

**Figure S10:** Principal component analysis of all the climatic, topographic and soil variables in Table S7.

##### 1.3.3 Correlation among the potential drivers

| | A | D | EH1[20km] | EH2[20km] | EH1[1.6km] | EH2[1.6km] | SHM | invEMT |
|---|---|---|---|---|---|---|---|---|
| **A** | - | - | - | - | - | - | - | - |
| **D** | **0.914** | - | - | - | - | - | - | - |
| **EH1[20km]** | -0.071 | -0.013 | - | - | - | - | - | - |
| **EH2[20km]** | 0.197 | 0.349 | **0.778** | - | - | - | - | - |
| **EH1[1.6km]** | 0.375 | 0.400 | 0.561 | 0.698 | - | - | - | - |
| **EH2[1.6km]** | 0.420 | 0.565 | 0.333 | **0.799** | 0.686 | - | - | - |
| **SHM** | 0.381 | 0.536 | 0.335 | 0.541 | 0.275 | 0.519 | - | - |
| **invEMT** | -0.009 | 0.126 | 0.197 | 0.082 | 0.218 | 0.044 | 0.406 | - |

**Table S8:** Correlations among the potential underlying drivers of the within-population genetic variation. Correlations higher than 0.7 are in bold.

### 2 Model equation and priors

We modeled each trait $y_{bpcr}$ such as:

$$y_{bpcr} \sim \mathcal{N}(\mu_{bpc}, \sigma_r^2)$$

$$\mu_{bpc} = \beta_0 + B_b + P_p + C_{c(p)}$$

where $\beta_0$ is the global intercept, $B_b$ the block intercepts, $P_p$ the population intercepts, $C_{c(p)}$ the clone intercepts and $\sigma_r^2$ the residual variance.

The prior of $\beta_0$ was weakly informative and centered around the mean of the observed values for the trait under considered, as follows:

$$\beta_0 \sim \mathcal{N}(\mu_y, 2)$$

The population and block intercepts, $P_p$ and $B_b$ were considered normally-distributed with variances $\sigma_P^2$ and $\sigma_B^2$, such as:

$$\begin{bmatrix} B_b \\ P_p \end{bmatrix} \sim \mathcal{N}\left(0, \begin{bmatrix} \sigma_B^2 \\ \sigma_P^2 \end{bmatrix}\right)$$

The clone intercepts $C_{c(p)}$ were considered to follow some population-specific normal distributions, such as:

$$C_{c(p)} \sim \mathcal{N}(0, \sigma_{C_p}^2)$$

where $\sigma_{C_p}^2$ are the population-specific variances among clones.

To partition the total variance, we parameterize our model so that only the total variance, $\sigma_{tot}^2$ has a prior, such that:

$$\sigma_{tot}^2 = \sigma_r^2 + \sigma_B^2 + \overline{\sigma_{C_p}^2} + \sigma_P^2$$

$$\sigma_r = \sigma_{tot} \times \sqrt{(\pi_r)}$$

$$\sigma_B = \sigma_{tot} \times \sqrt{(\pi_B)}$$

$$\sigma_P = \sigma_{tot} \times \sqrt{(\pi_P)}$$

$$\overline{\sigma_{C_p}} = \sigma_{tot} \times \sqrt{(\pi_C)}$$

$$\sigma_{tot} \sim \mathcal{S}^*(0, 1, 3)$$

where $\overline{\sigma_{C_p}}$ and $\overline{\sigma_{C_p}^2}$ are the mean of the population-specific among-clones standard deviations ($\sigma_{C_p}$) and variances ($\sigma_{C_p}^2$), respectively, and $\sum_l^4 \pi_l = 1$ (using the `simplex` function in `Stan`).

The population-specific among-clones standard deviations $\sigma_{C_p}$ follow a log-normal distribution with mean $\overline{\sigma_{C_p}}$ and variance $\sigma_K^2$, such as:

$$\sigma_{C_p} \sim \mathcal{LN}\left(\ln(\overline{\sigma_{C_p}}) - \frac{\sigma_K^2}{2} + \beta_X X_p, \sigma_K^2\right)$$

$$\sigma_K \sim \exp(1)$$

with $X_p$ the potential driver considered and $\beta_x$ its associated coefficient.

Here we provide further explanation regarding the use of the log-normal distribution for $\sigma_{C_p}$ in the model:

$$\sigma_{C_p} \sim \mathcal{LN}(\mu, \sigma_K^2) \Leftrightarrow \ln(\sigma_{C_p}) \sim \mathcal{N}(\mu, \sigma_K^2) \text{ with } \mu \text{ the median of } \sigma_{C_p}$$

$$\text{By definition: } \mathbb{E}(\sigma_{C_p}) = \exp\left(\mu + \frac{\sigma_K^2}{2}\right)$$

$$\text{We want: } \mathbb{E}(\sigma_{C_p}) = \exp\left(\mu + \frac{\sigma_K^2}{2}\right) = \sigma_{tot} \times \sqrt{(}\pi_4)$$

$$\text{Therefore: } \mu = \ln\left(\sigma_{tot} \times \sqrt{(}\pi_4)\right) - \frac{\sigma_K^2}{2}$$

### 3 Model accuracy on simulated data

We simulated data based on the real experimental design of two traits (height in Portugal at 20-month old and height in Bordeaux at 25-month old), which means that there were the same number of blocks, populations, clones per population and trees per clone as in the real experimental design. We ran 100 simulations, which are summarized in the tables below:

| Parameter | True value | Mean standard error | Mean bias of the mean | Mean bias of the median | 80% conf. int. coverage | 95% conf. int. coverage |
|---|---|---|---|---|---|---|
| $\beta_X$ | 0.1 | 0.055 | -0.002 | -0.003 | 83 | 97 |
| $\sigma_K$ | 0.1 | 0.067 | 0.020 | 0.014 | 92 | 99 |

**Table S9:** Summary of the 100 model outputs ran on simulated data based on height in Bordeaux at 25-month old.

| Parameter | True value | Mean standard error | Mean bias of the mean | Mean bias of the median | 80% conf. int. coverage | 95% conf. int. coverage |
|---|---|---|---|---|---|---|
| $\beta_X$ | 0.1 | 0.052 | -0.005 | -0.005 | 78 | 96 |
| $\sigma_K$ | 0.1 | 0.065 | 0.015 | 0.009 | 95 | 98 |

**Table S10:** Summary of the 100 model outputs ran on simulated data based on height in Portugal at 20-month old.

### 4 $\beta_X$ interpretation

We have:

$$\sigma_{C_p} \sim \mathcal{LN}\left(\ln(\overline{\sigma_{C_p}}) - \frac{\sigma_K^2}{2} + \beta_X \tilde{X}_p, \sigma_K^2\right)$$

with $\tilde{X}_p = (X_p - \mu_{X_p})/\sigma_{X_p}$ (the explanatory variables were scaled before the analyses; $\mu_{X_p}$ is the mean of $X_p$ and $\sigma_{X_p}$ is its standard deviation).

By definition:

$$\ln(\sigma_{C_p}) \sim \mathcal{N}\left(\ln(\overline{\sigma_{C_p}}) - \frac{\sigma_K^2}{2} + \beta_X \tilde{X}_p, \sigma_K^2\right)$$

We want to calculate the percent of change in $\sigma_{C_p}$ associated with a one-unit increase in $\tilde{X}_p$, that is a one-standard deviation increase in $X_p$. For that, we call $\sigma_{new}$ the value of $\sigma_{C_p}$ after increasing $\tilde{X}_p$ by one unit, and we have:

$$\begin{aligned}\ln(\sigma_{new}) &= \ln(\overline{\sigma_{C_p}}) - \frac{\sigma_K^2}{2} + \beta_X(\tilde{X}_p + 1) \\ &= \ln(\sigma_{C_p}) + \beta_X\end{aligned}$$

Therefore:

$$\begin{aligned}\ln(\sigma_{new}) - \ln(\sigma_{C_p}) &= \beta_X \\ \frac{\sigma_{new}}{\sigma_{C_p}} &= \exp(\beta_X) \\ 100 \times \left(\frac{\sigma_{new}}{\sigma_{C_p}} - 1\right) &= 100 \times (\exp(\beta_X) - 1) \\ 100 \times \left(\frac{\sigma_{new} - \sigma_{C_p}}{\sigma_{C_p}}\right) &= 100 \times (\exp(\beta_X) - 1)\end{aligned}$$

is the percent change in $\sigma_{C_p}$ associated with a one-unit increase in $\tilde{X}_p$ (that is, a one-standard deviation increase in $X_p$). For instance, a one-standard deviation increase in the inverse of the extreme minimum temperature is associated, on average, with $100 \times (\exp(-0.395) - 1) = -32.6\%$ change in $\sigma_{C_p}$ for height in Portugal, with $100 \times (\exp(-0.243) - 1) = -21.6\%$ change in $\sigma_{C_p}$ for height in Bordeaux at 25-month old and with $100 \times (\exp(-0.197) - 1) = -17.9\%$ change in $\sigma_{C_p}$ for height in Asturias. Similarly, a one-standard deviation increase in the summer heat moisture index is associated, on average, with $100 \times (\exp(-0.17) - 1) = -15.6\%$ change in $\sigma_{C_p}$ for height in Bordeaux at 25-month old and with $100 \times (\exp(-0.272) - 1) = -23.8\%$ change in $\sigma_{C_p}$ for height in Asturias.

### 5 Model outputs

#### 5.1 $\mathcal{R}^2$ estimates

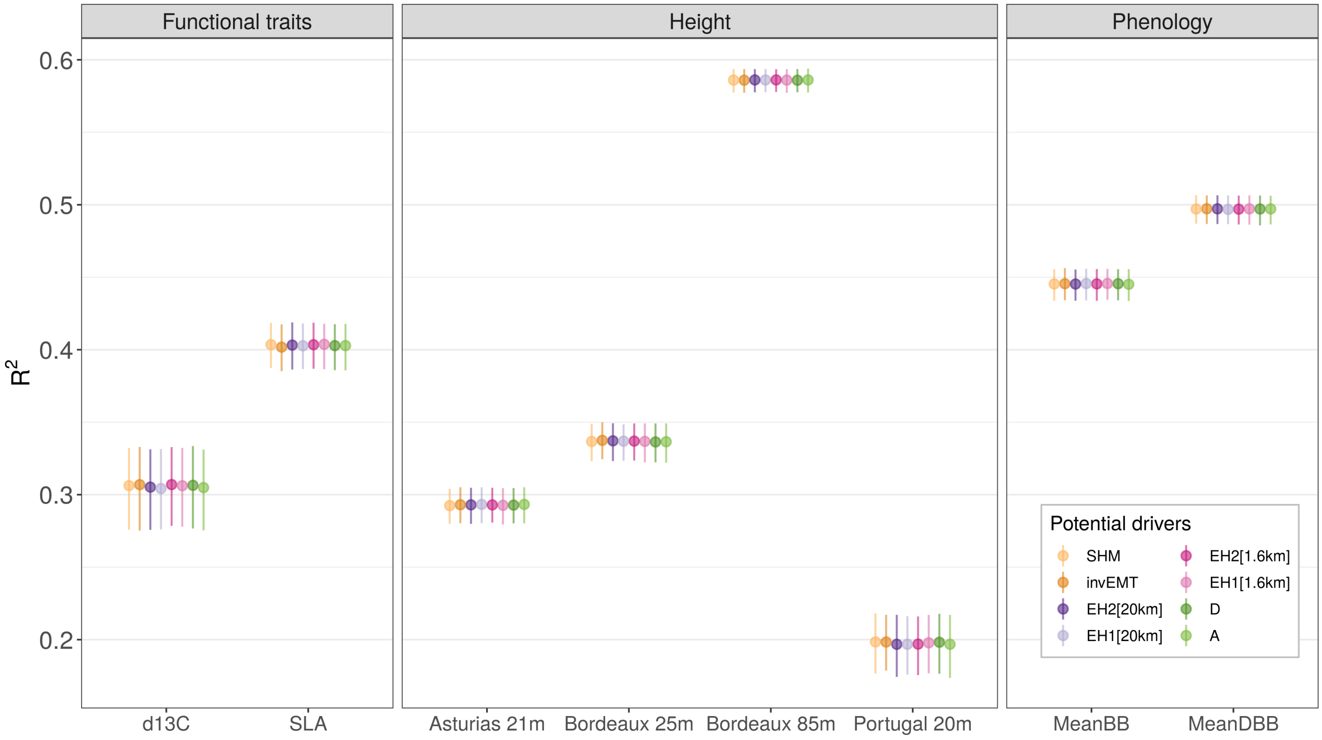

**Figure S11:** Median and 95% intervals of the posterior distributions of the total variance explained by the models ($\mathcal{R}^2$).

#### 5.2 $\beta_X$ estimates for the eight potential drivers

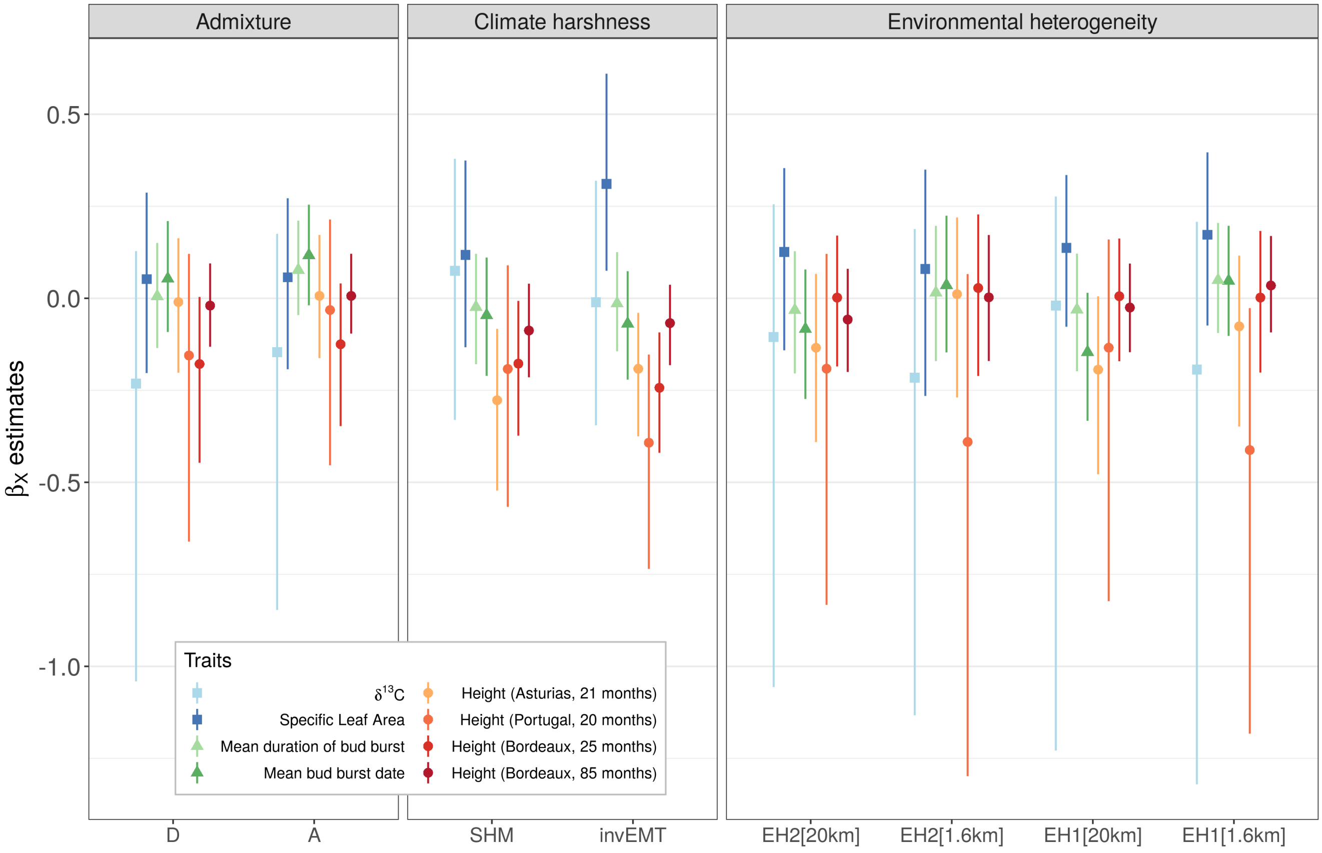

**Figure S12:** Median and 95% intervals of the posterior distributions of $\beta_X$, the coefficient corresponding to the potential drivers of the within-population genetic variation.

#### 5.3 $\sigma_{C_p}$ estimates and variance partitioning

##### 5.3.1 Height (Portugal, 20 months)

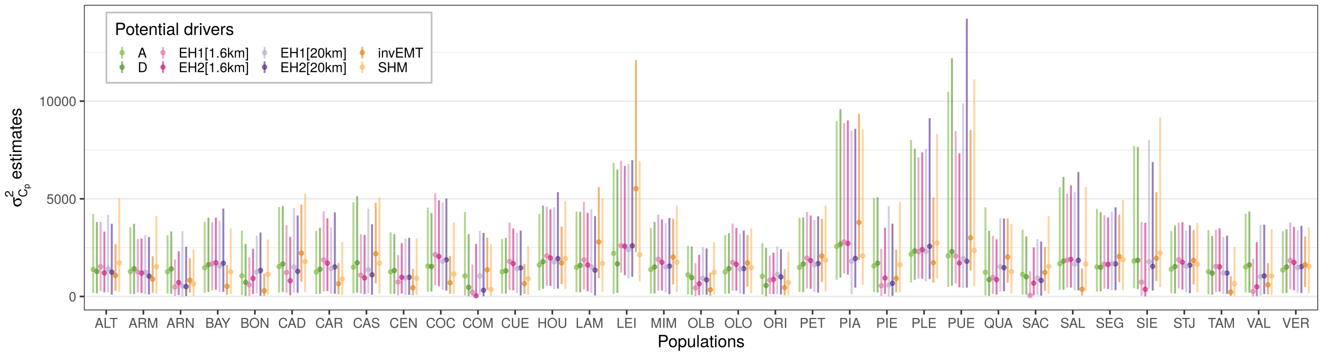

**Figure S13:** Median and 95% intervals of the posterior distributions of $\sigma^2_{C_p}$, corresponding to the within-population total genetic variance (i.e. population-specific among clones variance).

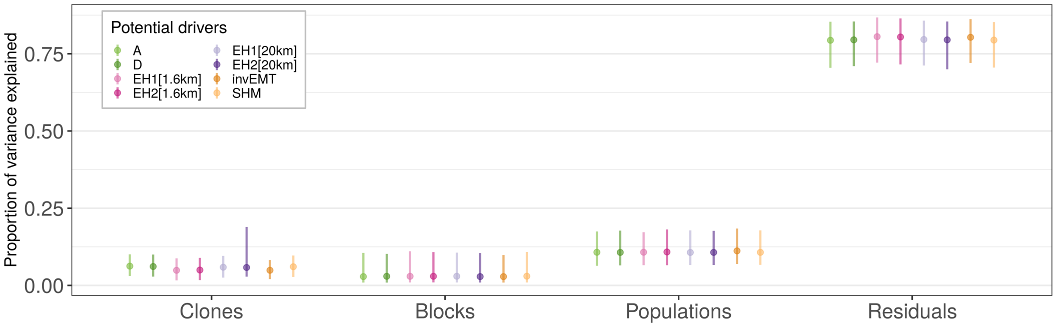

**Figure S14:** Proportion of variance explained by the different components, namely the clones ($\pi_C$), the blocks ($\pi_B$), the populations ($\pi_P$) and the residuals ($\pi_r$).

##### 5.3.2 Height (Bordeaux, 25 months)

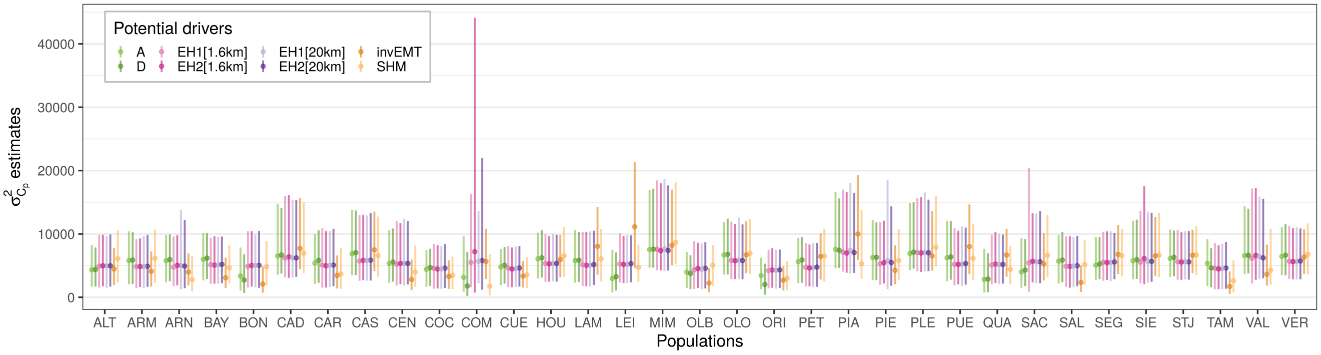

**Figure S15:** Median and 95% intervals of the posterior distributions of $\sigma^2_{C_p}$, corresponding to the within-population total genetic variance (i.e. population-specific among clones variance).

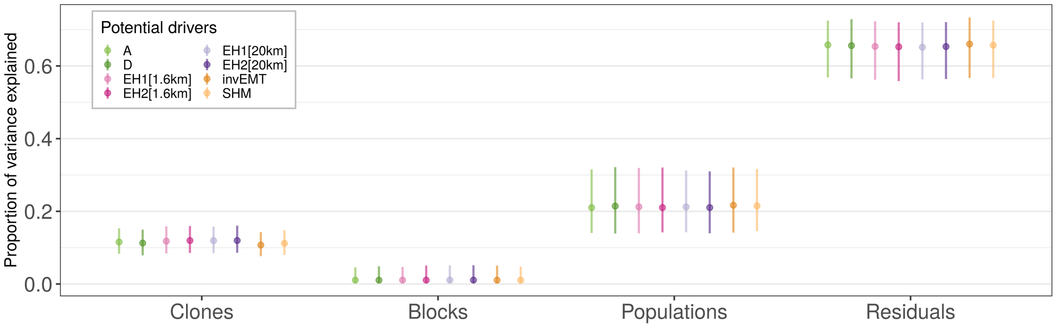

**Figure S16:** Proportion of variance explained by the different components, namely the clones ($\pi_C$), the blocks ($\pi_B$), the populations ($\pi_P$) and the residuals ($\pi_r$).

##### 5.3.3 Height (Bordeaux, 85 months)

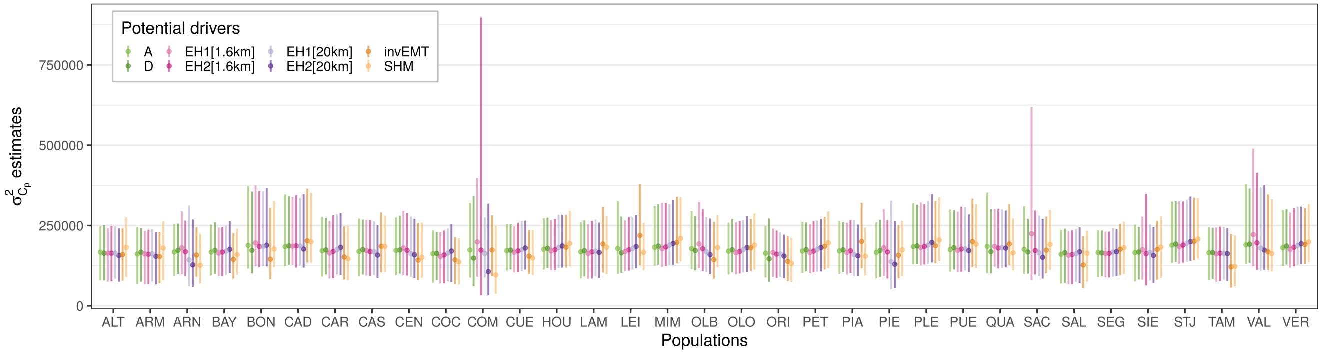

**Figure S17:** Median and 95% intervals of the posterior distributions of $\sigma^2_{C_p}$, corresponding to the within-population total genetic variance (i.e. population-specific among clones variance).

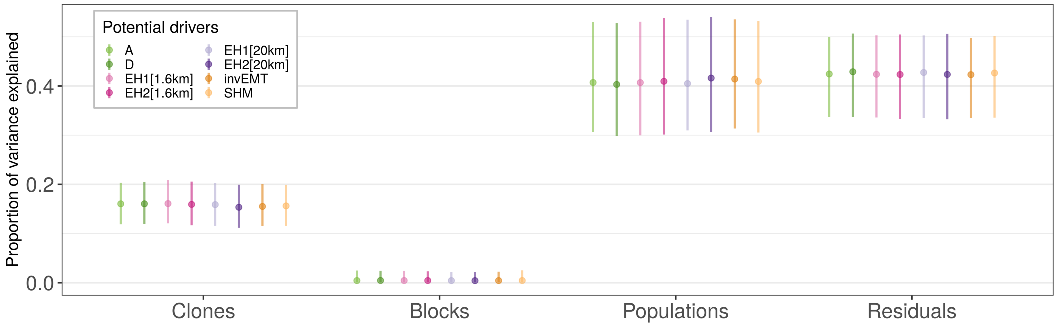

**Figure S18:** Proportion of variance explained by the different components, namely the clones ($\pi_C$), the blocks ($\pi_B$), the populations ($\pi_P$) and the residuals ($\pi_r$).

##### 5.3.4 Height (Asturias, 21 months)

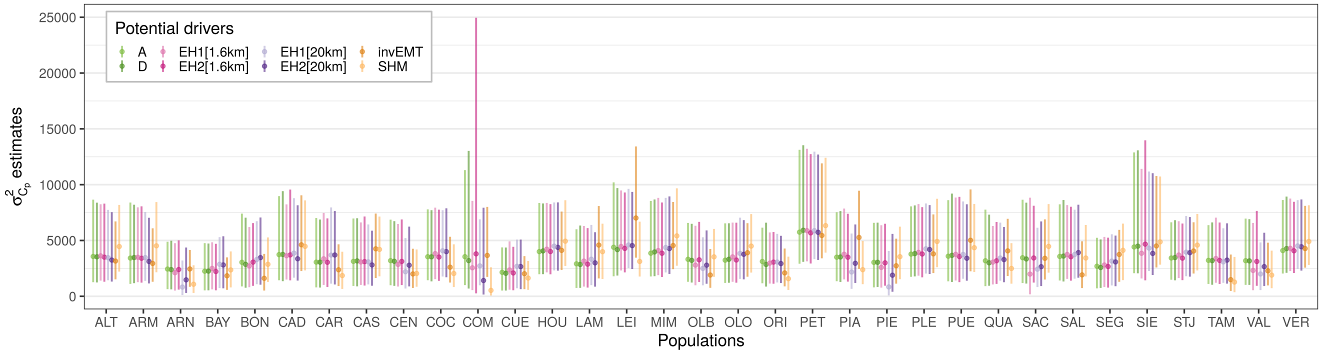

**Figure S19:** Median and 95% intervals of the posterior distributions of $\sigma^2_{C_p}$, corresponding to the within-population total genetic variance (i.e. population-specific among clones variance).

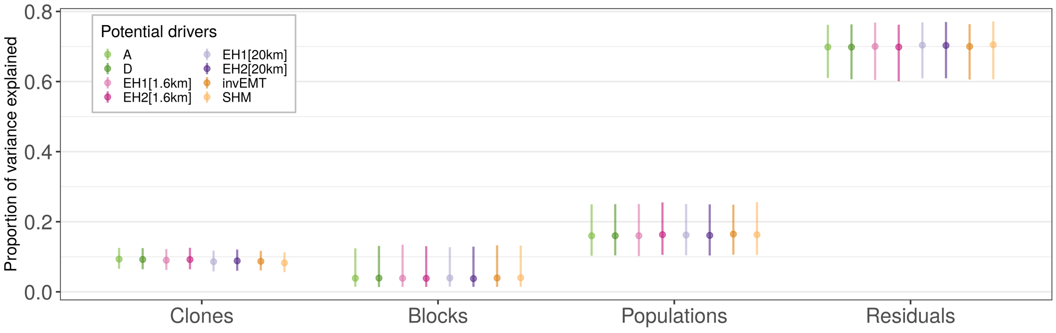

**Figure S20:** Proportion of variance explained by the different components, namely the clones ($\pi_C$), the blocks ($\pi_B$), the populations ($\pi_P$) and the residuals ($\pi_r$).

##### 5.3.5 Mean bud burst date (Bordeaux)

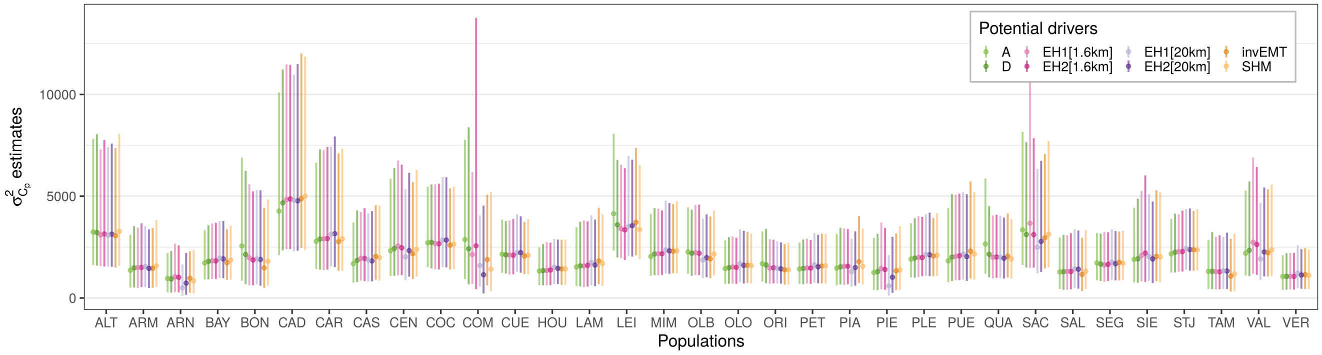

**Figure S21:** Median and 95% intervals of the posterior distributions of $\sigma^2_{C_p}$, corresponding to the within-population total genetic variance (i.e. population-specific among clones variance).

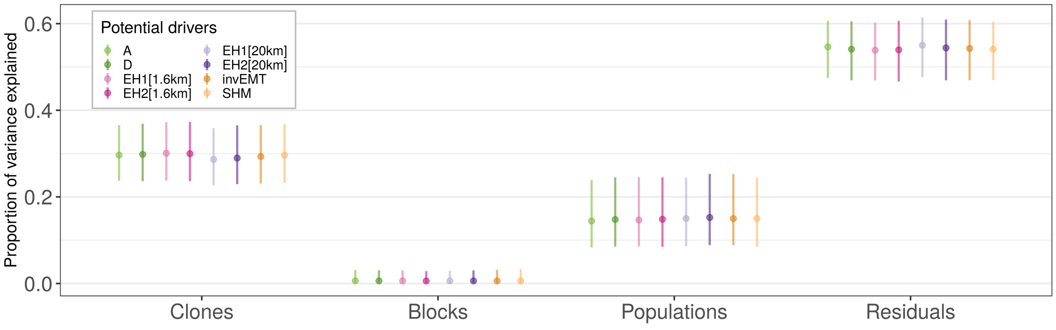

**Figure S22:** Proportion of variance explained by the different components, namely the clones ($\pi_C$), the blocks ($\pi_B$), the populations ($\pi_P$) and the residuals ($\pi_r$).

##### 5.3.6 Mean duration of bud burst (Bordeaux)

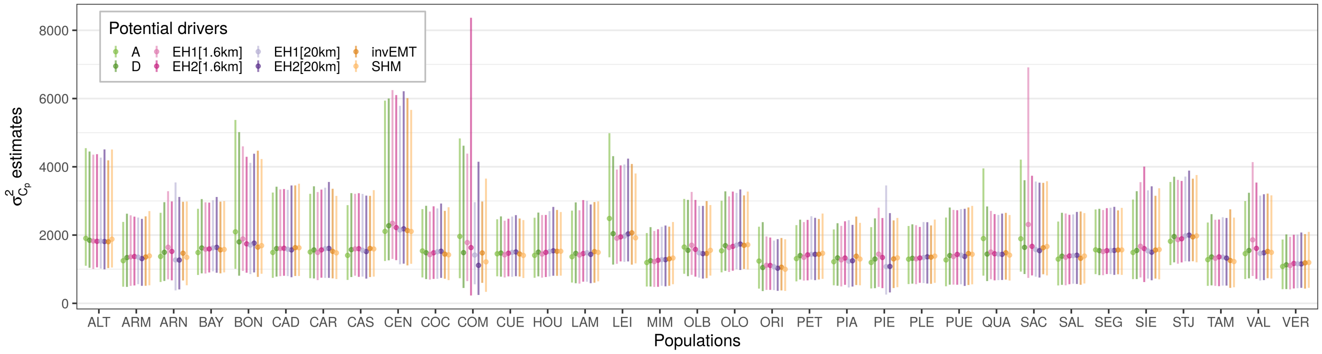

**Figure S23:** Median and 95% intervals of the posterior distributions of $\sigma^2_{C_p}$, corresponding to the within-population total genetic variance (i.e. population-specific among clones variance).

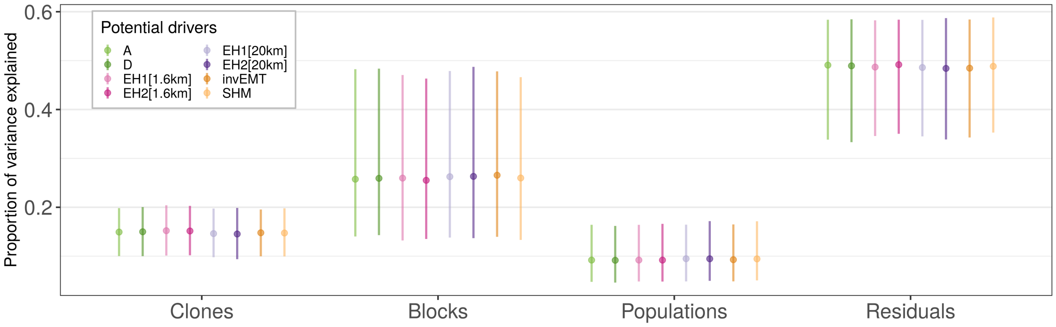

**Figure S24:** Proportion of variance explained by the different components, namely the clones ($\pi_C$), the blocks ($\pi_B$), the populations ($\pi_P$) and the residuals ($\pi_r$).

##### 5.3.7 Specific Leaf Area (Portugal)

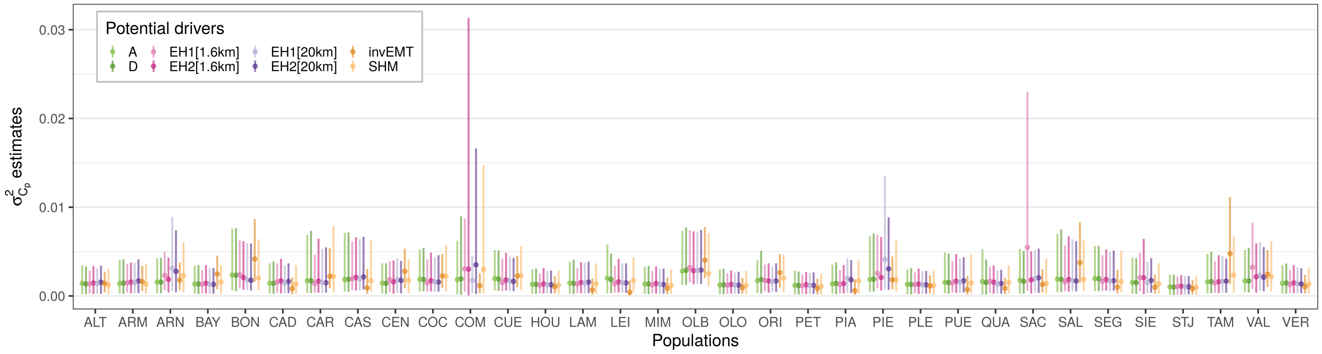

**Figure S25:** Median and 95% intervals of the posterior distributions of $\sigma^2_{C_p}$, corresponding to the within-population total genetic variance (i.e. population-specific among clones variance).

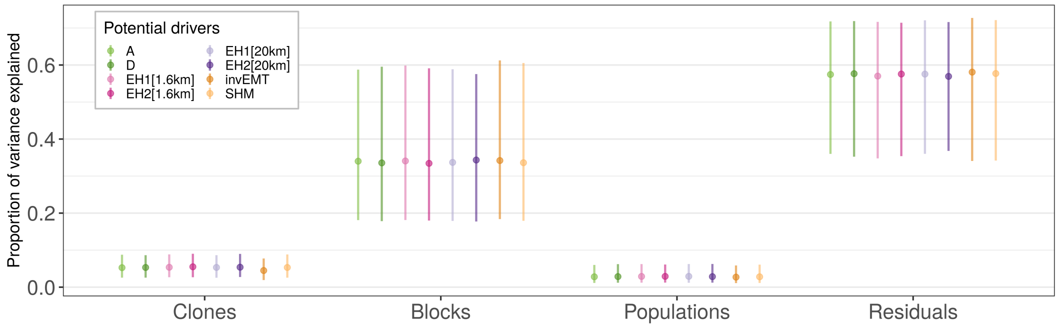

**Figure S26:** Proportion of variance explained by the different components, namely the clones ($\pi_C$), the blocks ($\pi_B$), the populations ($\pi_P$) and the residuals ($\pi_r$).

##### 5.3.8 $\delta^{13}$C (Portugal)

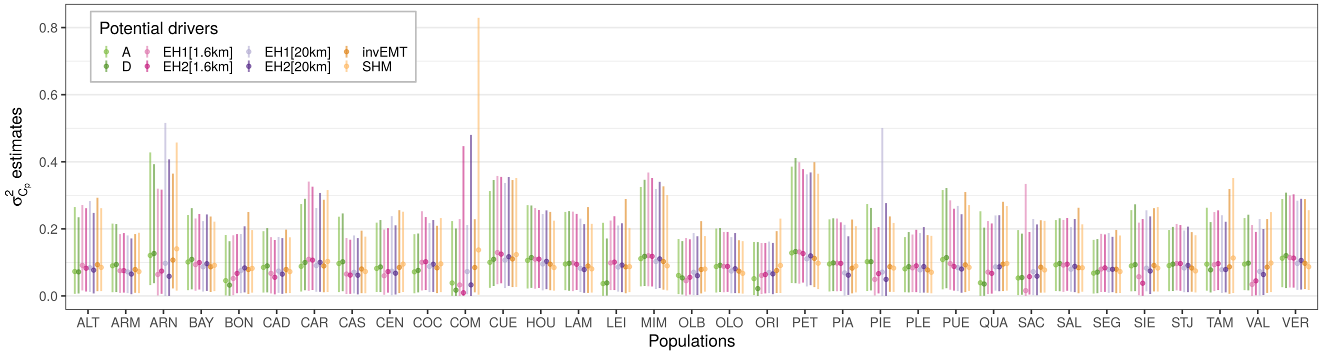

**Figure S27:** Median and 95% intervals of the posterior distributions of $\sigma^2_{C_p}$, corresponding to the within-population total genetic variance (i.e. population-specific among clones variance).

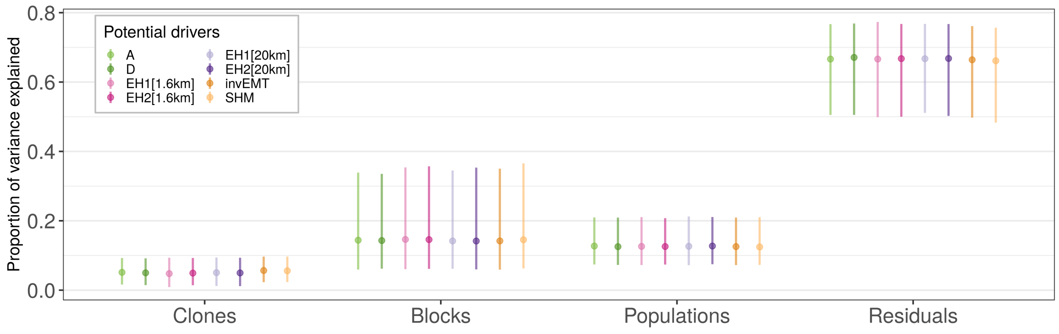

**Figure S28:** Proportion of variance explained by the different components, namely the clones ($\pi_C$), the blocks ($\pi_B$), the populations ($\pi_P$) and the residuals ($\pi_r$).

#### 5.4 Correlation between the number of clones per population and $\sigma_{C_p}$

| Drivers | Ht (Portugal) | Ht (Bordeaux 2013) | Ht (Bordeaux 2018) | Ht (Asturias) | meanBB | meanDBB | SLA | $\delta^{13}C$ |
|---|---|---|---|---|---|---|---|---|
| A | 0.035 | 0.047 | 0.119 | 0.105 | -0.217 | -0.161 | -0.119 | 0.152 |
| D | 0.038 | 0.068 | 0.149 | 0.122 | -0.238 | -0.189 | -0.107 | 0.147 |
| EH1[1.6km] | 0.423 | 0.024 | -0.283 | 0.317 | -0.276 | -0.381 | -0.364 | 0.519 |
| EH2[1.6km] | 0.480 | -0.166 | 0.087 | 0.066 | -0.285 | -0.262 | -0.399 | 0.564 |
| EH1[20km] | 0.272 | -0.042 | 0.350 | 0.321 | -0.080 | -0.093 | -0.245 | 0.440 |
| EH2[20km] | 0.404 | -0.035 | 0.497 | 0.433 | -0.110 | -0.006 | -0.424 | 0.567 |
| invEMT | 0.109 | 0.185 | 0.230 | 0.126 | -0.153 | -0.131 | -0.086 | 0.198 |
| SHM | 0.080 | 0.156 | 0.203 | 0.162 | -0.186 | -0.115 | -0.208 | -0.042 |

**Table S11:** Pearson correlation coefficients between the estimates of the within-population genetic variation (i.e. $\sigma_{C_p}$) and the number of clones per population, for each combination of trait and potential driver of the within-population genetic variation. The eight phenotypic traits are (from left to right): height in Portugal (October 2012), height in Bordeaux (France, November 2013), height in Bordeaux (France, November 2018), height in Asturias (Spain, November 2012), mean bud burst date in Bordeaux (over the years 2013, 2014, 2015 and 2017), mean duration of bud burst in Bordeaux (over the years 2014, 2015 and 2017), specific leaf area in Portugal and the isotope discrimination of $\delta^{13}C$ in Portugal.

### 6 Climatic transfer distances

We estimated the association between climatic transfer distances for both EMT and SHM and the within-population genetic variation. The climatic transfer distances were calculated as the absolute difference between the climate in the location of origin of each population and the climate in the test site, for instance the climatic transfer distance for EMT between the population p and the common garden s was: equal to $abs(CTD_{EMT,s,p})$ with $CTD_{EMT,s,p} = EMT_p - EMT_s$.

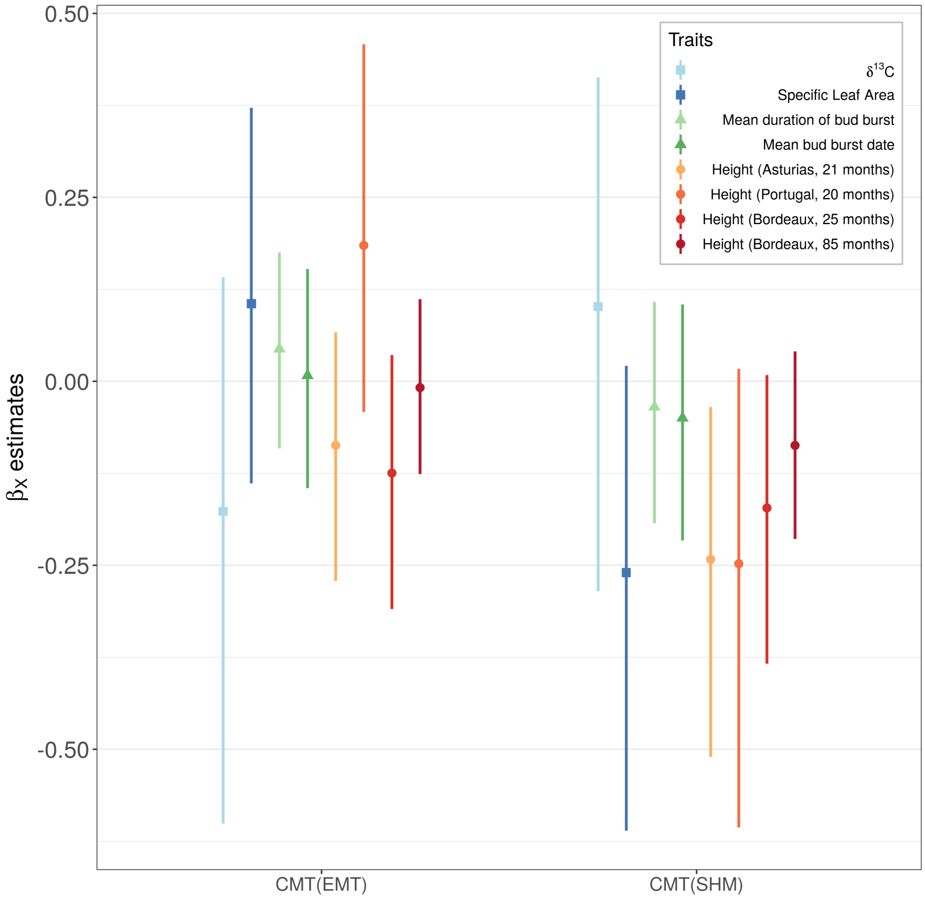

**Figure S29:** Median and 95% intervals of the posterior distributions of $\beta_X$, the coefficient corresponding to the potential drivers (here the climatic transfer distances) of the within-population genetic variation.

We did not detect an association between climatic transfer distance and within-population genetic variation for SLA, as hypothesized in the discussion.

### 7 Validation step

In the validation step, we performed exactly the same analyses (i.e. using the same model formula and code) as for the CLONAPIN height data, but on independent height data kindly provided by Ricardo Alia. This independent height data comes from a progeny test near Asturias, planted in 2005, and in which 23 provenances are shared with the CLONAPIN data (see Tables S12 and S13).

#### 7.1 Experimental design and exploratory analyses

| Populations | Mean | Variance | Number of families | Number of individuals |
|---:|---:|---:|---:|---:|
| **ALT** | 116.42373 | 1,697.8691 | 7 | 59 |
| **ARM** | 127.13793 | 2,275.1203 | 8 | 87 |
| **ARN** | 115.44615 | 1,723.3808 | 12 | 130 |
| **BAY** | 99.32948 | 1,503.3269 | 16 | 173 |
| **CAD** | 132.54000 | 2,112.1499 | 10 | 100 |
| **CAR** | 92.02128 | 1,415.1952 | 4 | 47 |
| **CAS** | 110.46970 | 2,974.8683 | 6 | 66 |
| **CEN** | 108.35000 | 1,003.4641 | 4 | 40 |
| **COC** | 109.85714 | 1,847.6504 | 8 | 77 |
| **CUE** | 98.58389 | 908.6635 | 14 | 149 |
| **LAM** | 118.09459 | 2,725.6211 | 7 | 74 |
| **LEI** | 114.00862 | 3,988.9825 | 12 | 116 |
| **MIM** | 115.20000 | 1,388.7236 | 9 | 90 |
| **ORI** | 91.91156 | 1,777.4784 | 15 | 147 |
| **PIA** | 123.42553 | 1,373.9454 | 4 | 47 |
| **PIE** | 106.27778 | 828.5654 | 2 | 18 |
| **PLE** | 122.73077 | 2,050.5288 | 9 | 104 |
| **PUE** | 119.90000 | 1,735.9694 | 5 | 50 |
| **SAL** | 97.21250 | 1,034.1695 | 8 | 80 |
| **SEG** | 117.18354 | 1,917.2846 | 16 | 158 |
| **SIE** | 116.38596 | 2,180.2412 | 5 | 57 |
| **TAM** | 86.90728 | 1,139.1780 | 16 | 151 |
| **VAL** | 106.48352 | 1,632.6747 | 8 | 91 |

**Table S12:** Mean, variance, number of families and number of individuals in each population for the height measurements in the progeny test near Asturias when the trees were 3-year old.

| Populations | Mean | Variance | Number of families | Number of individuals |
|---:|---:|---:|---:|---:|
| **ALT** | 343.4138 | 14,256.949 | 7 | 58 |
| **ARM** | 346.8605 | 15,262.263 | 8 | 86 |
| **ARN** | 315.4656 | 8,490.866 | 12 | 131 |
| **BAY** | 275.3810 | 9,417.399 | 16 | 168 |
| **CAD** | 372.6667 | 10,699.909 | 10 | 96 |
| **CAR** | 255.5778 | 8,827.659 | 4 | 45 |
| **CAS** | 312.1746 | 16,498.792 | 6 | 63 |
| **CEN** | 297.4250 | 9,311.892 | 4 | 40 |
| **COC** | 308.0789 | 9,849.060 | 8 | 76 |
| **CUE** | 277.6667 | 8,059.922 | 14 | 147 |
| **LAM** | 318.2676 | 16,332.885 | 7 | 71 |
| **LEI** | 331.9057 | 18,746.258 | 12 | 106 |
| **MIM** | 327.0444 | 8,190.200 | 9 | 90 |
| **ORI** | 262.3333 | 9,773.510 | 15 | 144 |
| **PIA** | 349.7234 | 6,867.813 | 4 | 47 |
| **PIE** | 295.5000 | 5,594.618 | 2 | 18 |
| **PLE** | 356.3010 | 11,389.154 | 9 | 103 |
| **PUE** | 336.8269 | 11,972.773 | 5 | 52 |
| **SAL** | 280.4872 | 6,440.773 | 8 | 78 |
| **SEG** | 338.6382 | 11,798.934 | 16 | 152 |
| **SIE** | 330.2500 | 12,265.718 | 5 | 56 |
| **TAM** | 272.7919 | 6,791.801 | 16 | 149 |
| **VAL** | 293.1000 | 10,550.788 | 8 | 90 |

**Table S13:** Mean, variance, number of families and number of individuals in each population for the height measurements in the progeny test near Asturias when the trees were 6-year old.

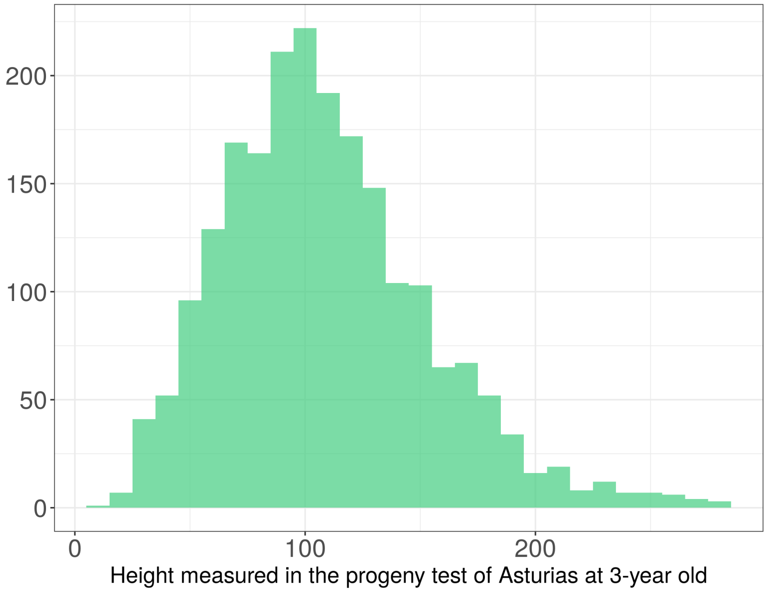

**Figure S30:** Distribution of height measurements at 3-year old in the progeny test near Asturias (independent dataset used for the validation analysis).

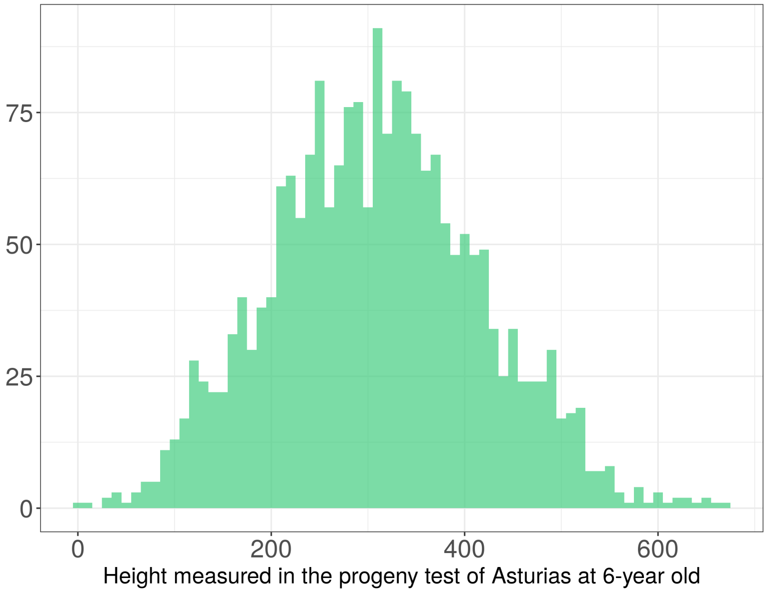

**Figure S31:** Distribution of height measurements at 6-year old in the progeny test near Asturias (independent dataset used for the validation analysis).

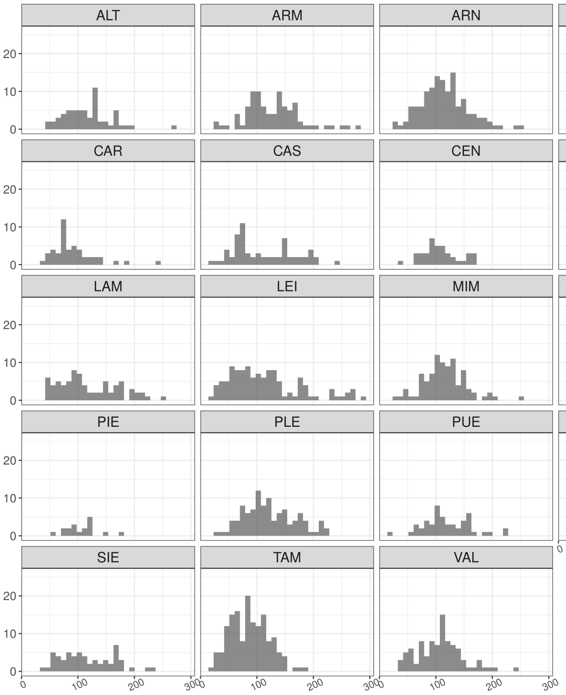

**Figure S32:** Height distribution at 3-year old for the 23 populations shared between the CLONAPIN dataset and the independent dataset used in the validation analysis, i.e. a progeny test near Asturias.

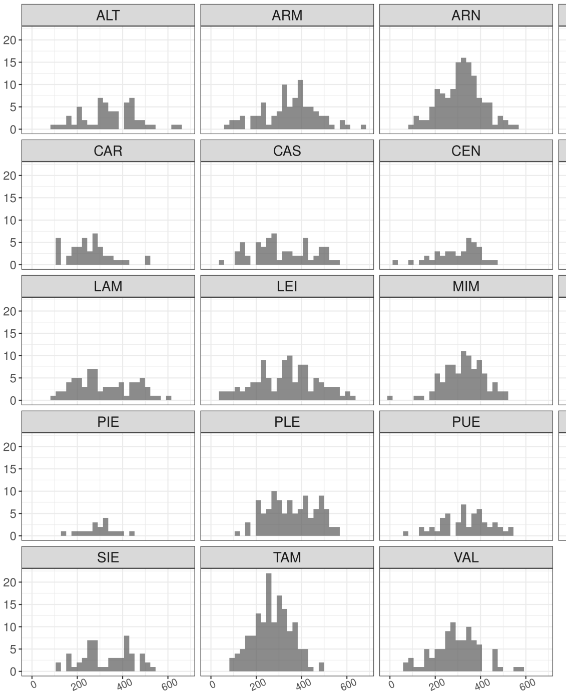

**Figure S33:** Height distribution at 6-year old for the 23 populations shared between the CLONAPIN dataset and the independent dataset used in the validation analysis, i.e. a progeny test near Asturias.

#### 7.2 Model equation and priors

We used the same mathematical model as the one used on CLONAPIN data (see section 2 in the Supplementary Information) but replacing clones by families.

We modeled each trait $y_{bpfr}$ such as:

$$\begin{aligned} y_{bpfr} &\sim \mathcal{N}(\mu_{bpcf}, \sigma_r^2) \\ \mu_{bpf} &= \beta_0 + B_b + P_p + F_{f(p)} \end{aligned} \tag{1}$$

where $\beta_0$ is the global intercept, $B_b$ the block intercepts, $P_p$ the population intercepts, $F_{f(p)}$ the family intercepts and $\sigma_r^2$ the residual variance.

The prior of $\beta_0$ was weakly informative and centered around the mean of the observed values for the trait under considered, as follows:

$$\beta_0 \sim \mathcal{N}(\mu_y, 2)$$

The population and block intercepts, $P_p$ and $B_b$ were considered normally-distributed with variances $\sigma_P^2$ and $\sigma_B^2$, such as:

$$\begin{bmatrix} B_b \\ P_p \end{bmatrix} \sim \mathcal{N}\left(0, \begin{bmatrix} \sigma_B^2 \\ \sigma_P^2 \end{bmatrix}\right)$$

The family intercepts $F_{f(p)}$ were considered to follow some population-specific normal distributions, such as:

$$F_{f(p)} \sim \mathcal{N}(0, \sigma_{F_p}^2)$$

where $\sigma_{F_p}^2$ are the population-specific variances among families.

To partition the total variance, we parameterize our model so that only the total variance, $\sigma_{tot}^2$ has a prior, such that:

$$\begin{aligned} \sigma_{tot}^2 &= \sigma_r^2 + \sigma_B^2 + \overline{\sigma_{F_p}^2} + \sigma_P^2 \\ \sigma_r &= \sigma_{tot} \times \sqrt{(\pi_r)} \\ \sigma_B &= \sigma_{tot} \times \sqrt{(\pi_B)} \\ \sigma_P &= \sigma_{tot} \times \sqrt{(\pi_P)} \\ \overline{\sigma_{F_p}} &= \sigma_{tot} \times \sqrt{(\pi_F)} \\ \sigma_{tot} &\sim \mathcal{S}^*(0, 1, 3) \end{aligned} \tag{2}$$

where $\overline{\sigma_{F_p}}$ and $\overline{\sigma_{F_p}^2}$ are the mean of the population-specific among-families standard deviations ($\sigma_{F_p}$) and variances ($\sigma_{F_p}^2$), respectively, and $\sum_l^4 \pi_l = 1$ (using the `simplex` function in `Stan`).

The population-specific among-families standard deviations $\sigma_{F_p}$ follow a log-normal distribution with mean $\overline{\sigma_{F_p}}$ and variance $\sigma_K^2$, such as:

$$\sigma_{F_p} \sim \mathcal{LN}\left(\ln(\overline{\sigma_{F_p}}) - \frac{\sigma_K^2}{2} + \beta_X X_p, \sigma_K^2\right)$$
$$\sigma_K \sim \exp(1) \quad (3)$$

with $X_p$ the potential driver considered and $\beta_x$ its associated coefficient.

##### 7.3 $\beta_X$ estimates for the eight potential drivers

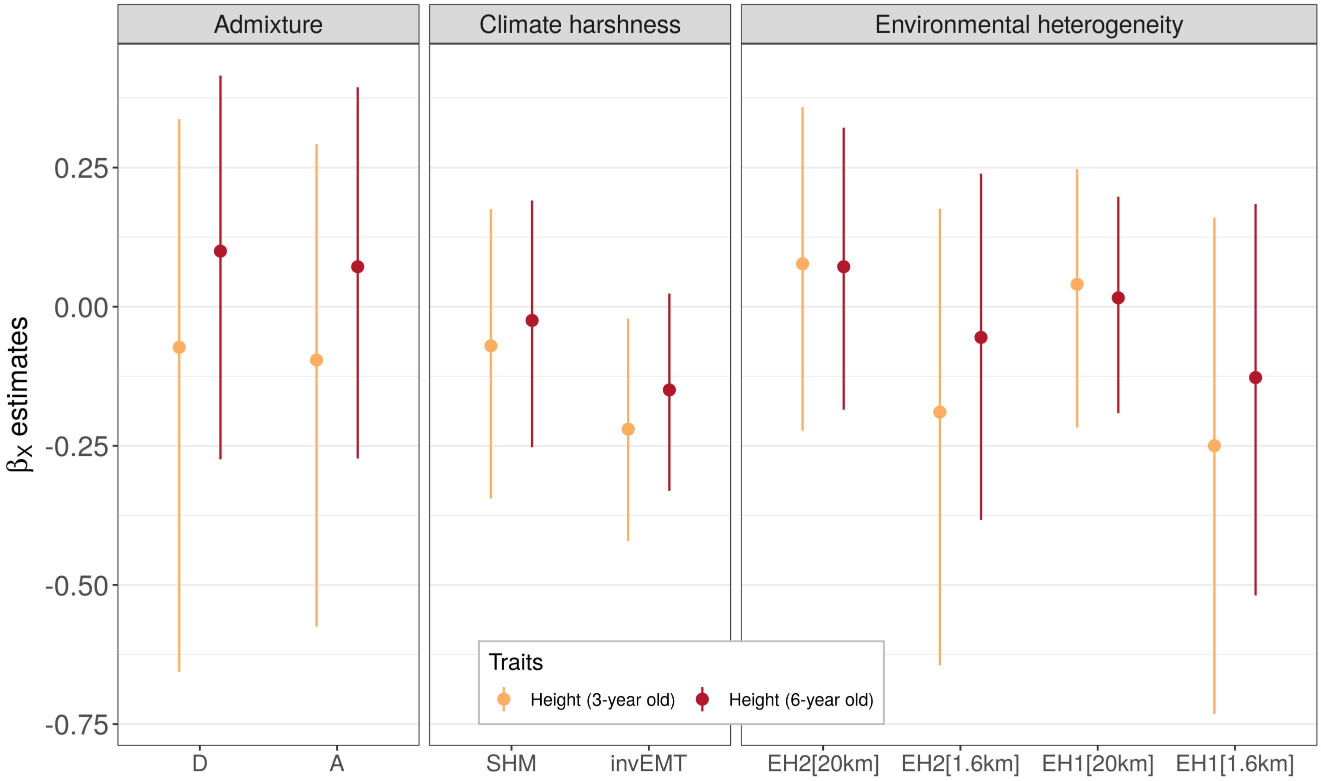

**Figure S34:** Median and 95% intervals of the posterior distributions of $\beta_X$, the coefficient corresponding to the association between the eight potential drivers and the within-population additive genetic variation.

#### 7.4 $\sigma_{C_p}$ estimates and variance partitioning

##### 7.4.1 Height at 3-year old

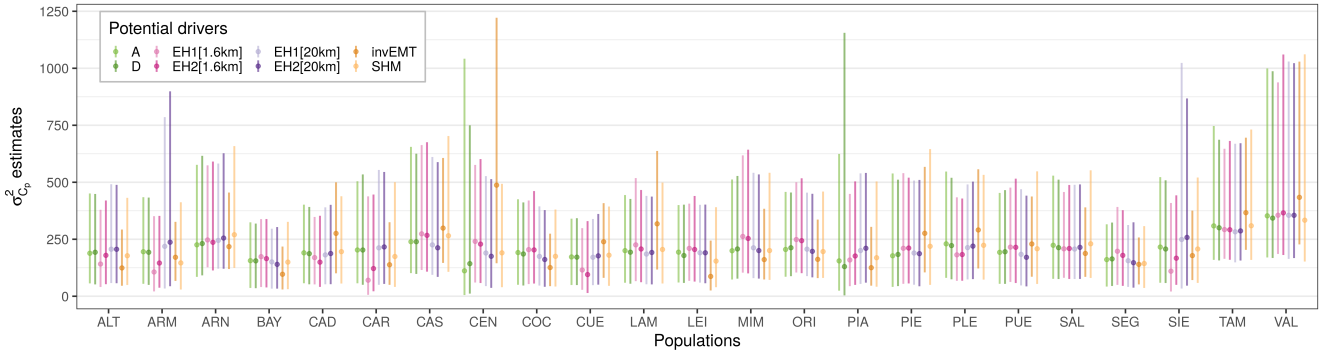

**Figure S35:** Median and 95% intervals of the posterior distributions of $\sigma^2_{C_p}$, corresponding to the association between the within-population additive genetic variance (i.e. population-specific among families variance) and the eight potential drivers.

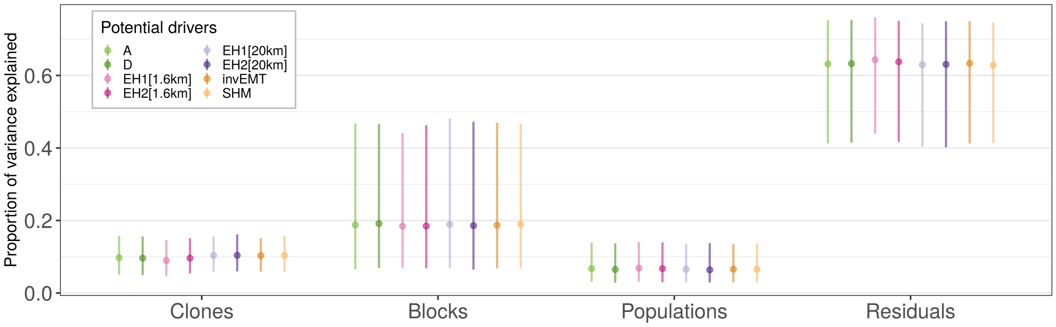

**Figure S36:** Proportion of variance explained by the different components, namely the clones ($\pi_C$), the blocks ($\pi_B$), the populations ($\pi_P$) and the residuals ($\pi_r$).

##### 7.4.2 Height at 6-year old

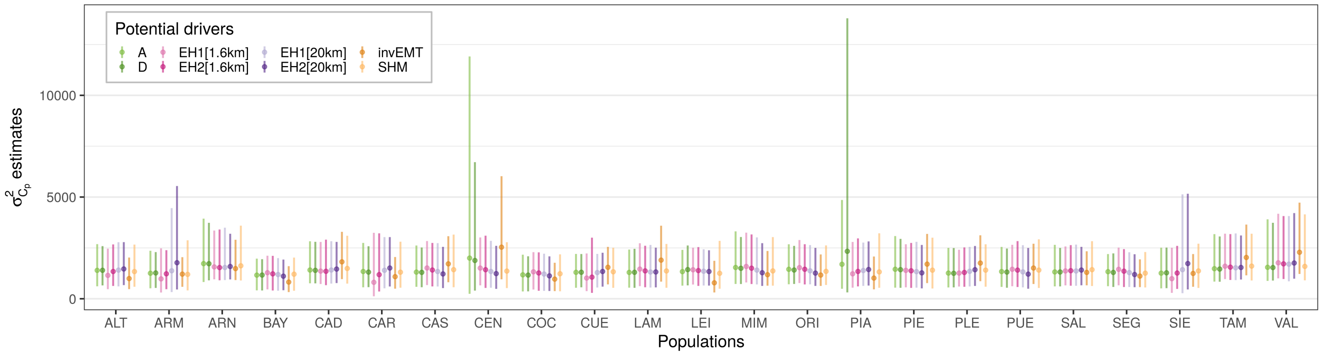

**Figure S37:** Median and 95% intervals of the posterior distributions of $\sigma^2_{C_p}$, corresponding to the within-population total genetic variance (i.e. population-specific among clones variance).

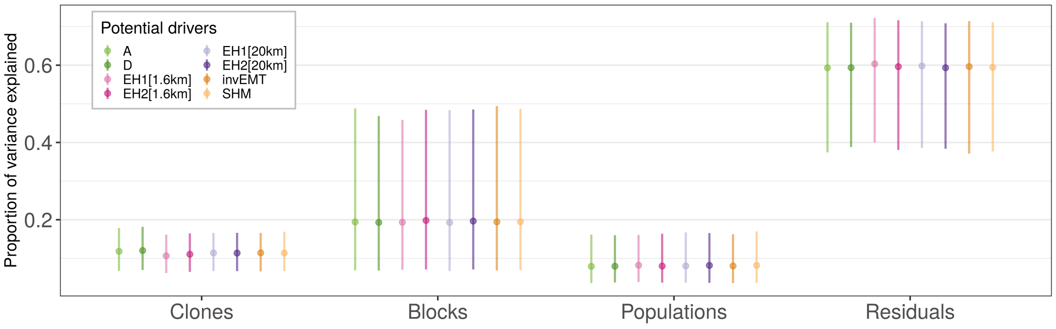

**Figure S38:** Proportion of variance explained by the different components, namely the clones ($\pi_C$), the blocks ($\pi_B$), the populations ($\pi_P$) and the residuals ($\pi_r$).

### 8 Changes since preregistration

This study was pre-registered at the Center for Open Science (https://osf.io/knx6z/?view_only=41bb7b5cbf72 41d0856e8b9e393cc795). Some changes have been made in the final manuscript compared to what was indicated in the pre-registration. There are listed below:

- There was a mistake in Table 2 of the pre-registration: we did not have the soil moisture index (SMI). Moreover, the notation for the summer heat moisture index has changed: instead of SumHMI, it is now noted as SHM.

- The initial number of clones and populations were 523 and 34 respectively. However, calculating the genetic variation in one population (from Madisouka) was impossible as there was only one clone in that population. That's why there are 522 clones and 33 populations in the final manuscript.

- In the part *Statistical models* of the pre-registration, we specified: 'We will test three different models, from the simplest to the most complex, and we will keep the complex model if it converges and if the credible intervals are not too wide compared to the simpler models.' As the most complex worked well, we did not run the simpler models and we directly used the most complex model in the manuscript. In addition, there was a mistake in the pre-registration formula regarding the estimation of $\sigma_{C_P}$ with the log-normal distribution, which was corrected in the final manuscript (see section *Model equation and priors* of the Supplementary Information).

- In the part *EnvironmentIndices* of the pre-registration, we indicated that we would calculate EHW, the environmental heterogeneity in a 20-km around each population location, as the variance of the PC1 scores weighted by the relative probability of gene flow from the surrounding region. However, the pollen dispersal kernels we used (from de-Lucas et al. 2008) are highly leptokurtic, which means that the probability of gene flow among trees located more than 200m from the GPS coordinates of the population is very low. Since the resolution of the climatic variables was only 1 × 1 km, we obtained implausible values for EHW and therefore decided not to use it. Instead, we calculated the variance of PC1 scores within a 1.6 km radius of the population locations. Furthermore, as the first two components of the PCA both explained a large part of the environmental variation (45.2% and 34.1%, respectively; S10), we decided to calculate the environmental heterogeneity indices based on the PC1 and PC2 scores, resulting in four indices in the end: EH1[20km], EH2[20km], EH1[1.6km] and EH2[1.6km]. Last, we indicated that we would use the soil moisture index (SMI) as a measure of climate harshness but that was a mistake, we used the summer heat moisture index (SHM).
